# Structures of the Otopetrin Proton Channels Otop1 and Otop3

**DOI:** 10.1101/542308

**Authors:** Kei Saotome, Bochuan Teng, Che Chun (Alex) Tsui, Wen-Hsin Lee, Yu-Hsiang Tu, Mark S. P. Sansom, Emily R. Liman, Andrew B. Ward

## Abstract

Otopetrins (Otop1-Otop3) comprise one of only two known eukaryotic proton-selective channel families. Otop1 is required for formation of otoconia and is a candidate mammalian sour taste receptor. Here, we report cryo-EM structures of zebrafish Otop1 and chicken Otop3 in lipid nanodiscs. The structures reveal a dimeric architecture of Otopetrins with each subunit consisting of twelve transmembrane helices divided into structurally related N and C domains. Cholesterol-like molecules occupy various sites in Otop1 and Otop3 and occlude a cavernous central tunnel. Two hydrophilic vestibules, as well as the intrasubunit interface between N and C domains, form conduits for water entry into the membrane plane in molecular dynamics simulations, suggesting they each could provide pathways for proton conduction. We also demonstrate the functional relevance of a salt bridge in the C domain vestibule by mutagenesis. Our results provide a structural basis for understanding the function of the Otopetrin proton channel family.

## Main Text

Proton channels mediate the passage of protons across cell membranes, thereby regulating the cellular and extracellular pH as well as membrane potential^1^. The diverse biological roles of proton channel activity include the triggering of bioluminescence in dinoflagellates^2^, regulation of pH in lung epithelia^3^, and the detection of sour taste^4-6^. Knowledge of proton channel physiology and molecular mechanisms is largely derived from, and limited to, the M2 proton channel of influenza^7^ and the eukaryotic voltage-gated Hv1 proton channel^8,9^. Recently, Otopetrins were identified as a novel family of eukaryotic proton channels^10^. Mice have three related genes (Otop1, Otop2 and Otop3) that encode channels with distinct biophysical properties. For example, the current amplitudes of mouse Otop1 and mouse Otop3 increase linearly as a function of extracellular pH over a range of pH 6 – pH 4, while the current of mouse Otop2 saturates at ∼pH 5. Otop1 has been demonstrated to be proton-selective, with a remarkable 2 × 10^5^-fold selectivity for protons over Na^+10^.

Because they were only recently characterized, the physiological roles of Otopetrins are just now beginning to be uncovered. Notably, Otop1 is required for proton currents in murine cells that detect sour taste and is a likely candidate for a sour taste receptor^4-6,10^. In addition, prior genetic studies identified Otop1 as the gene mutated in a spontaneously occurring vestibular disorder in mice and required in the vestibular system for normal otoconia development in mice^11^ and zebrafish^12,13^. Otop1 is also highly expressed in brown and white adipose tissue and plays a role in insulin resistance^14^. Otop2 and Otop3 have been detected in the digestive tract and elsewhere in the body^10,15^ suggesting yet unappreciated roles for proton conduction in various cell types.

The recent characterization of Otopetrins as proton channels has opened avenues to decipher their physiological, biophysical, and biochemical characteristics. Otopetrins are not related to M2, Hv1, or other known ion channels, indicating they operate using novel structure-function principles. To provide a starting point for mechanistic studies on Otopetrins, we conducted cryoelectron microscopy (cryo-EM) studies of zebrafish Otop1 and chicken Otop3, which are 30% identical to each other by sequence and share 44% and 59% identity with human OTOP1 and OTOP3, respectively (Supplementary Fig. 1). When expressed in HEK-293 cells, both zebrafish Otop1 and chicken Otop3 conduct proton currents in response to lowering the extracellular pH like their mammalian counterparts (Fig. 1a, e, Supplementary Fig. 2)^10^. For the remainder of this manuscript we refer to these proteins as Otop1 and Otop3, respectively. We purified and reconstituted full-length proteins in lipid nanodiscs and determined C2-symmetric structures at overall resolutions of 3.0 Å for Otop1 and 3.3 Å for Otop3 (Fig. 1, Supplementary Figs. 3, 4, Supplementary Table 1). The quality of the cryo-EM maps was sufficient to define side chain orientation of almost all amino acids in the transmembrane helices (Supplementary Fig. 5). Notably, there are numerous lipid densities surrounding the channels or embedded in an internal lipid tunnel at the dimer interface (Fig. 1). These lipids were either carried over from cells or added during purification and nanodisc reconstitution. Of these, two densities adopted distinctive shapes and were modeled as cholesterol while an additional six densities corresponded to cholesteryl hemisuccinate (CHS) in the Otop1 map, whereas two CHS molecules were built into the Otop3 map (Supplementary Fig. 5c-f).

**Figure 1.**
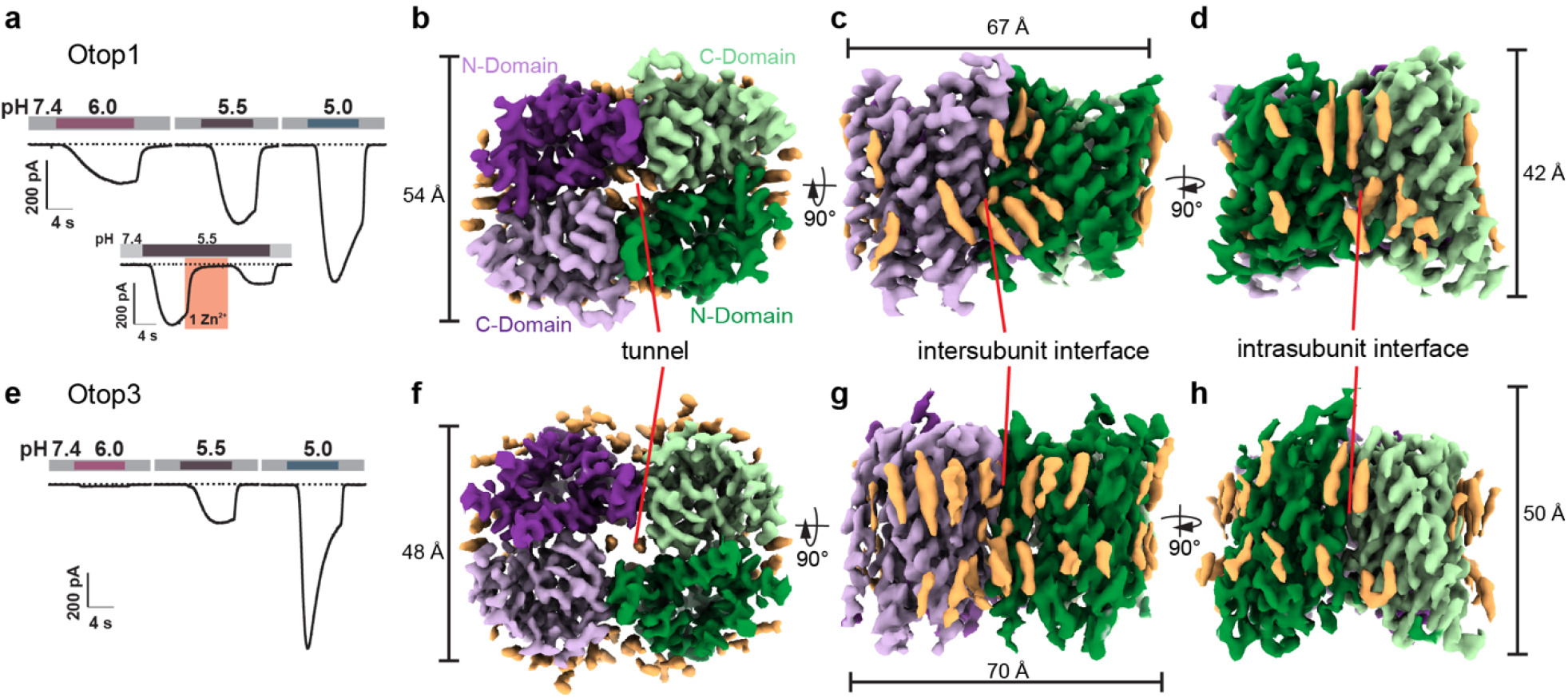
Proton channel function and structures of Otop1 and Otop3 homodimers in lipid nanodiscs. **a**, currents measured by whole-cell patch clamp recording in a HEK-293 cell expressing zfOtop1 in response to acidic extracellular solutions with the indicated pH (pH_i_ = 7.4, V_m_ = −80 mV). Inset shows inhibition of Otop1 currents by 1 mM Zn^2+^ (pink bar). **b**-**d,** orthogonal views of sharpened cryo-EM maps of zfOtop1. One subunit is colored dark and light shades of purple, while the other subunit is colored green and light green. Each subunit has a structurally homologous N domain (dark shade) and C domain (light shade). The center of the dimer contains a tunnel occupied by cholesterol-like densities. Putative cholesterol and lipid densities are colored light orange. **e,** currents measured by whole cell patch clamp recording in a HEK-293 cell expressing chOtop3 in response to acidic extracellular solutions with pH indicated (pH_i_= 7.4, V_m_ = −80mV). **f-h**, orthogonal views of sharpened cryo-EM maps of chOtop3, colored as in (**b-d**).

**Figure 2.**
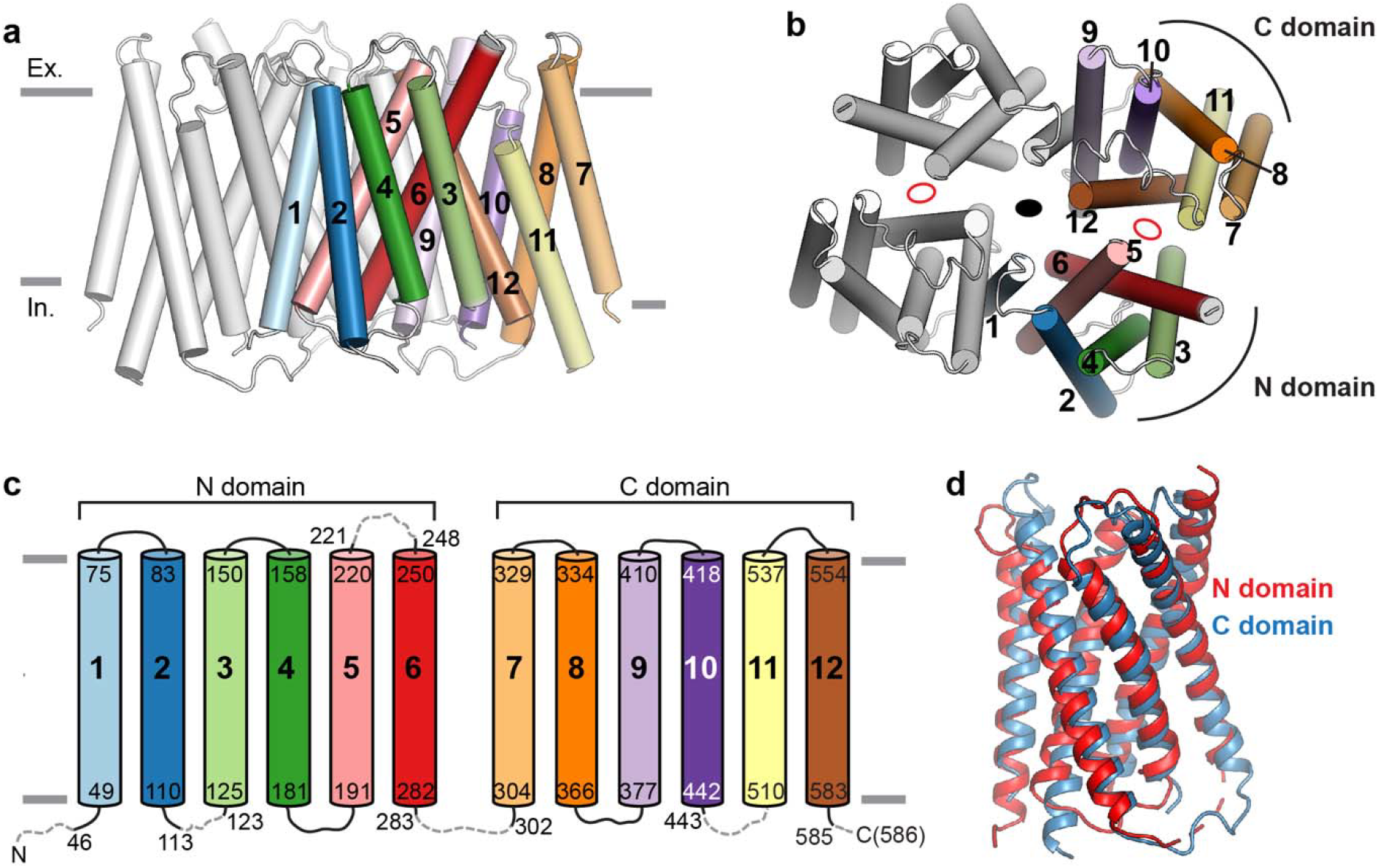
Domain organization and conservation. **a,b**, side (**a**) and top (**b**) views of model of Otop1 dimer showing TM helices as cylinders. One subunit is colored gray while in the other subunit each of the twelve TM helices are colored differently and numbered. In **b**, the central two-fold symmetry axis is depicted as a black oval, and pseudosymmetry axes relating the N domain and C domain within each subunit are depicted as empty red ovals. **c,** schematic of Otop1 structure, with TM helices colored corresponding to **a, b**. Loops not modeled due to poor density are depicted as dashed gray lines. **d,** structural alignment of the N domain (red) and C domain (blue).

**Figure 3.**
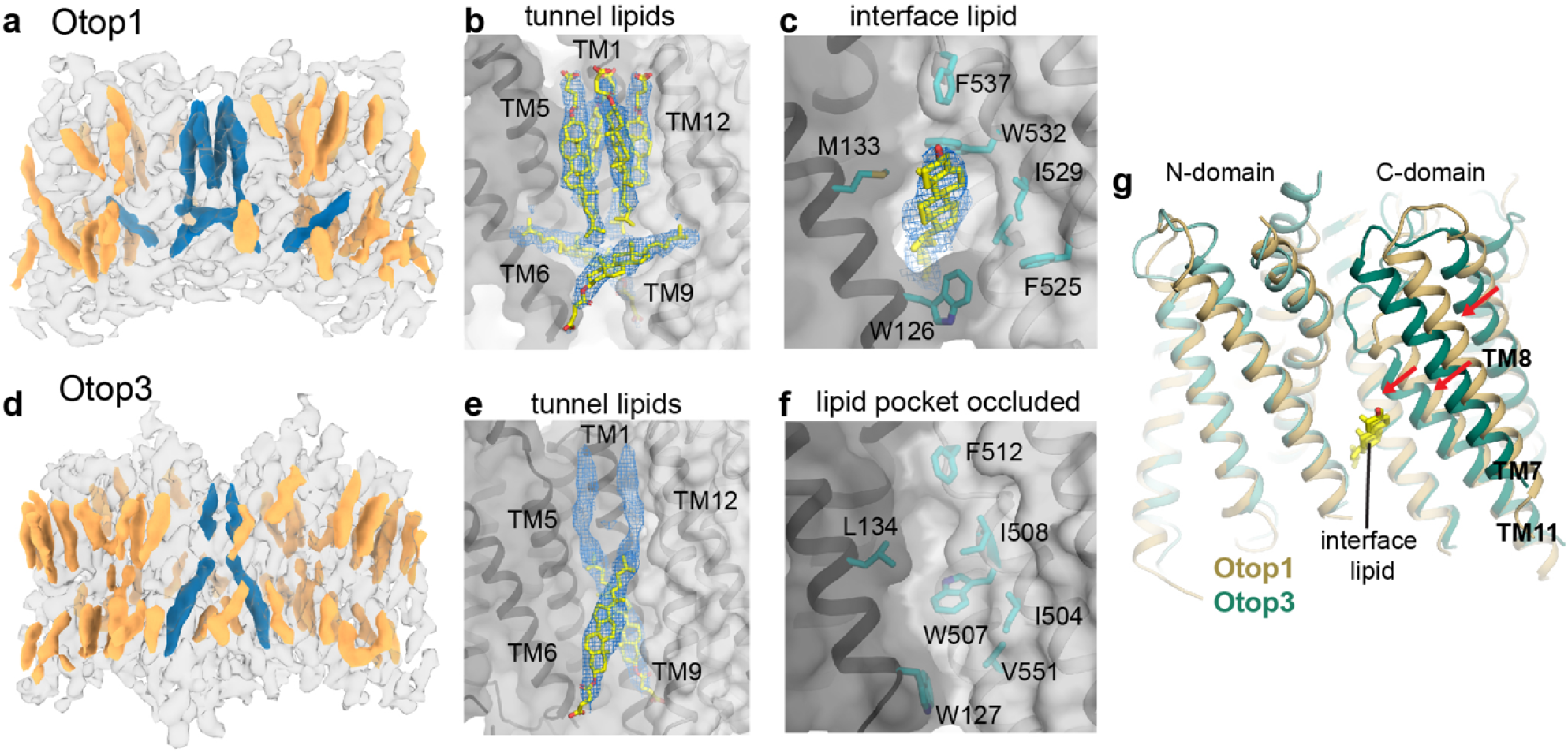
Lipid binding in Otop1 and Otop3. **a, d,** side views of cryo-EM map of Otop1 (**a**) and Otop3 (**d**). Protein density is transparent gray, annular lipid densities are colored gold, and lipid/cholesterol densities at the N domain-C domain intrasubunit interface and lipid tunnel are colored dark blue. **b**, densities (blue mesh, 3σ) corresponding to three CHS molecules (yellow and red sticks) in the lipid tunnel are apparent in Otop1**. c,** Density (blue mesh, 3σ) corresponding to a cholesterol molecule is also present at the intrasubunit interface. **e**, densities corresponding to a CHS molecule (blue mesh, 3σ) and an unassigned lipid (blue mesh, 3σ) are present in the Otop3 lipid tunnel. **f**, there is no lipid at the intrasubunit interface in Otop3, and the pocket is occluded. **g**, superimposition of Otop1 and Otop3 single subunits based on the N-domain shows that TM7, TM8, and TM11 of Otop3 is shifted relative to Otop1, likely due to absence of interface lipid.

**Figure 4.**
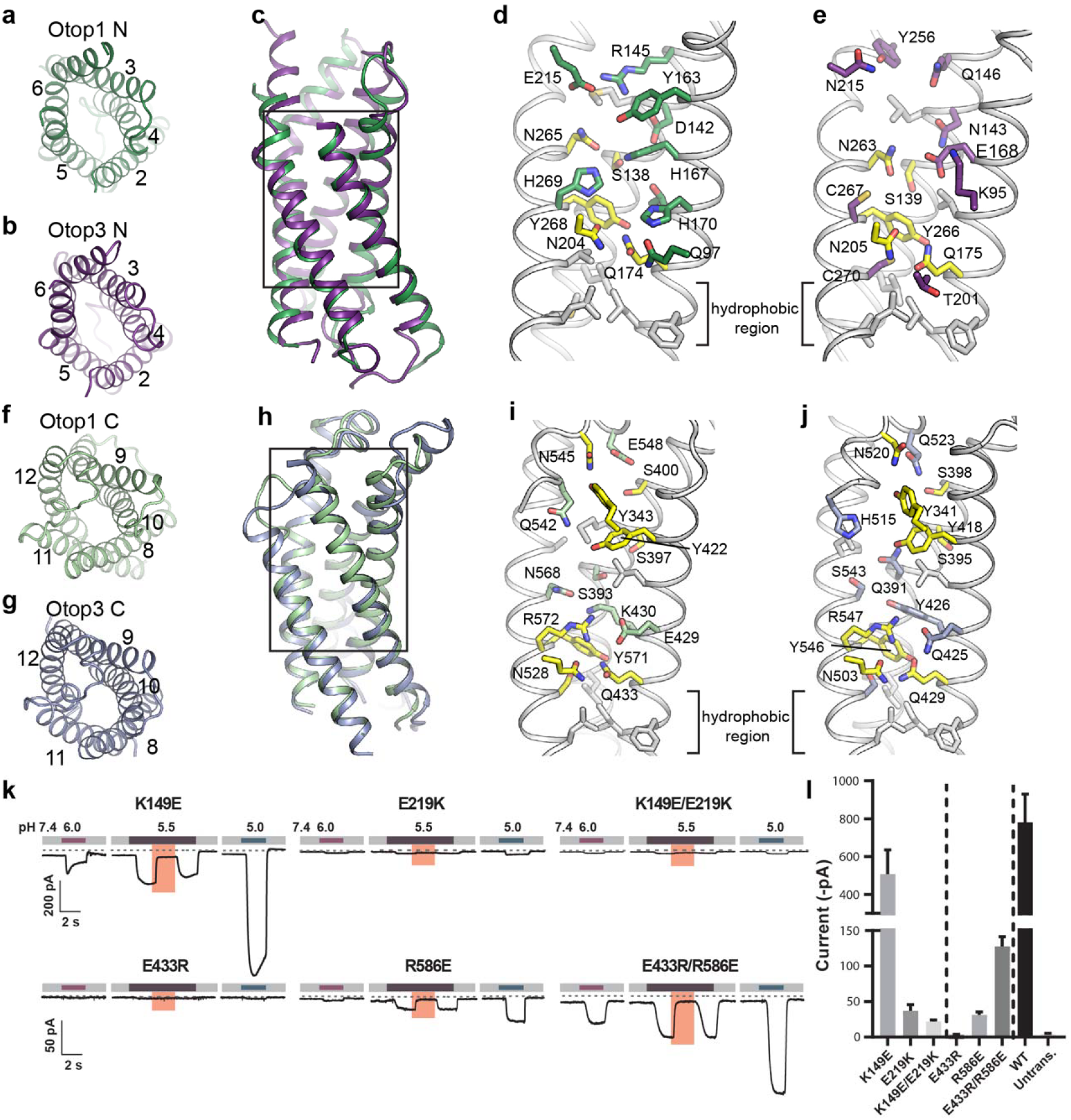
N and C domain vestibules Otop1 and Otop3. **a, b,** top views of Otop1 N domain (**a**) and Otop3 N domain (**b**). **c**, alignment of N-terminal domains of Otop1 (green) and Otop3 (purple). TM1 is not shown. **d, e,** expanded views of the boxed region in (**c**) for Otop1 (**d**) and Otop3 (**e**). **f, g,** top views of C domains of Otop1 (light green) and Otop3 (light purple). **h,** alignment of C domains of Otop1 (green) and Otop3 (purple). TM7 is not shown. **i, j,** expanded views of the boxed region in **d** for Otop1 (**e**) and Otop3 (**f**). In **d, e, i, j** residues conserved between Otop1 and Otop3 are colored yellow. **k**, currents measured by whole-cell patch clamp recording in HEK-293 cells expressing each mouse Otop1 mutant in response to acidic extracellular solutions with pH indicated (pH_i_ = 7.4, V_m_ = −80 mV). The currents elicited in response to pH 5.5 were inhibited by 1 mM Zn^2+^ (pink bars). Numbering is according to the mouse Otop1 sequence. **l**, averaged current magnitude in response to pH 5.0 from experiments as in the left panel. Error bars represent S.E.M.

**Figure 5.**
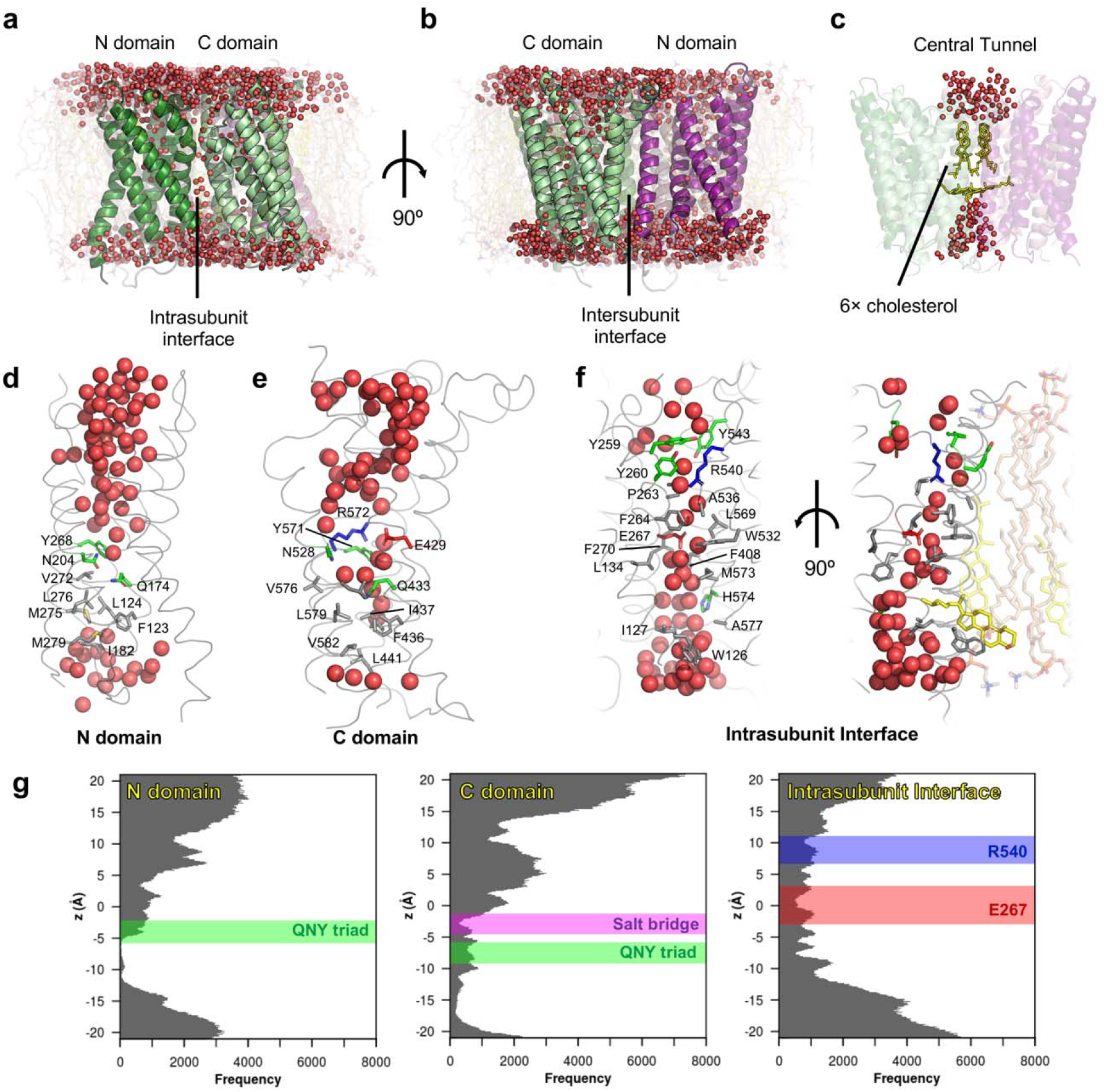
Molecular dynamics simulation reveals hydration of potential proton pathways in Otop1. **a-f**, snapshots at the end of a 100-ns all-atom simulation (Run 3 referring to Fig S10; **d-f** show snapshots from subunit 1). A continuous presence of water is observed at the intrasubunit interface (**a**) but not at the intersubunit interface (**b**). **c**, cholesterol molecules in the central tunnel exclude water passage completely. In **d, e,** the QNY triad in both N and C domains appear to separate the upper wetted and the less hydrated regions. In **f**, the water wire found at the intrasubunit interface is more uniform and stable than that in N and C domains. In the right-hand panel, the cholesterol molecule initially present at the interface is colored solid yellow, while other cholesterol and POPC molecules in the proximity are colored transparent yellow and wheat respectively. **g**, the distribution of water oxygen atoms along the three potential proton pathways, averaged across two subunits and three simulations. The region along the z-axis occupied by the fluctuations of relevant conserved side chain positions are drawn as rectangles, covering the 1^st^–9^th^ deciles range.

## Results

### Overall structure and domain organization

The structures show that Otop1 and Otop3 are cuboid-shaped homodimers with dimensions of roughly 70 Å × 50 Å × 50 Å, and almost all of their ordered mass resides within the membrane (Fig. 1). A single Otopetrin subunit can be divided into two halves, the N and C domains. In the homodimeric arrangement, the N and C domains from two subunits occupy four quadrants surrounding a central axis, resulting in a pseudotetrameric organization (Fig. 1b, f). A cavernous tunnel containing lipids and/or cholesterols coincides with the central two-fold axis of Otop1 and Otop3. Intersubunit and intrasubunit interfaces between N and C domains stabilize the overall assembly (Fig. 1c, d, g, h).

The topological organization of Otop1 (Fig. 2) is shared by Otop3 (Supplementary Fig. 6a), and therefore likely to be adopted among all Otopetrins. Each Otop1 subunit contains twelve transmembrane helices (TM1-TM12) grouped into two bundles, the N (TM1-6) and C (TM7-TM12) domains. The N and C domains are structurally similar and are related by a pseudo two-fold symmetry axis at the intrasubunit interface (Fig. 2). This domain organization is reminiscent of, but distinct from, the major facilitator superfamily (MFS) transporters^16,17^ and resistance-nodulation-cell division (RND) family of membrane proteins^18^ (Supplementary Fig. 6b-d). The intrasubunit interface of Otop1 is mediated mainly by TM3, TM5, TM6, and TM12, and is highly conserved, suggesting a critical role in structural and functional integrity (Fig. 2, Supplementary Fig. 7a-g). The interdimer interface, mediated by TM1 and TM9, is less conserved between Otop1 and Otop3 and across all Otopetrins (Supplementary Fig. 7a-c, h-l). We used fluorescence-detection size exclusion chromatography^19^ (FSEC) to test the role of specific residues on dimer formation and identified two alanine substitutions at two tryptophan residues (W394A and W398A) on TM9 of Otop1 that resulted in monomeric species (Supplementary Fig. 7j). These mutants confirm the importance of these residues in dimerization and suggest that dimerization is stabilized by burial of large hydrophobic residues at the interface.

### Lipids and cholesterols in Otop1 and Otop3

Facilitated by nanodisc reconstitution, numerous annular and bound lipids were captured and resolved in the structures of Otop1 and Otop3, revealing how the molecules interact with the membrane (Fig. 3, Supplementary Fig. 5c). Most remarkably, the central tunnel of Otop1 harbors six robust densities that correspond to CHS molecules (Fig. 3a, b, Supplementary Fig. 5d). We performed all-atom molecular dynamics (MD) simulations with cholesterol molecules placed in the CHS sites. The arrangement of cholesterol molecules in the tunnel is maintained throughout the (sub-microsecond) MD simulations, indicating that they represent stable binding poses for cholesterol-like moieties (Supplementary Fig. 8a–c). Otop3 has a distinct arrangement of lipids in the central tunnel, including two identifiable CHS molecules (Figs. 1, 3, Supplementary Fig. 5e,f). CHS, which was added during purification of Otop1 and Otop3, greatly enhances the monodispersity of detergent-solubilized proteins as shown by FSEC (Supplementary Figs. 3a, 4a). In native bilayers, cholesterol or cholesterol-like lipids likely bind in the central tunnel to stabilize the dimeric protein assembly^20,21^. Indeed, in simulations from which cholesterol was omitted, we observed considerable conformational drifts over the course of 100 ns (Supplementary Fig. 8d). The bound cholesterol molecules may also play a crucial functional role by occluding the relatively large tunnel, thereby preventing the flow of water, ions or other solutes.

At the intrasubunit interface of Otop1, a well-resolved cholesterol-like density is sandwiched in a pocket between the N and C domains in the lower leaflet (Fig. 3c). In contrast to those in the central tunnel, these cholesterols show greater mobility on this simulation timescale. In most cases, these cholesterol molecules reposition themselves during the simulation (Supplementary Fig. 8a-c). This lipid is absent in the Otop3 density map, despite high sequence conservation in this region (Supplementary Fig. 7a). Instead, the pocket is occluded by a conserved tryptophan (W507 in Otop3), resulting in a shift of the C-domain relative to the N-domain (Fig. 3g, h). Occupancy of this interface lipid site therefore appears to influence the conformation of Otopetrins. Overall, these observations underscore the importance of lipids, in particular cholesterol-like molecules, in shaping Otopetrin structure, and suggest that lipids play a central role in dimer stability, function, and/or membrane localization. The differences in lipid-binding observed when comparing Otop1 and Otop3 suggest that lipid modulation is subtype-specific.

### Putative pores in the N and C domains

Protons are believed to transfer across membrane proteins by ‘hopping’ along a hydrogen-bonded network consisting of water molecules and/or protein groups^1,22-26^. The proton conduction pathway(s) in Otopetrins would thus likely be at least partially hydrated. Two structurally analogous vestibule-shaped openings in each Otop1/Otop3 subunit could represent loci for proton permeation, one housed by the N domain and the other by the C domain (Fig. 4). The N domain vestibule is lined by TM2-TM6 (Fig. 4a-c), while the C domain vestibule is lined by TM8-TM12 (Fig. 4f-h). Both of the vestibules contain numerous polar and charged residues, many of which are conserved, apparently creating favorable environment for water molecules (Fig. 4d, e, i, j). Below the bilayer midpoint lies a region of hydrophobic residues that constrict the N and C domain vestibules in both Otop1 and Otop3, potentially serving as hydrophobic ‘plugs’ that regulate water or ion accessibility. Notably, hydrophobic gates are exploited across many ion channel families, including Hv1^27,28^.

In Hv1, an arginine-aspartic acid salt bridge has been proposed to function as a ‘selectivity filter’ in the proton conduction pathway^26,29^. Each of the vestibules of Otop1 also contains an apparent salt bridge, R145-E215 in the N domain vestibule (Fig. 4d) and E429-R572 in the C domain vestibule (Fig. 4i). To test the functional significance of these salt bridges, we introduced charge reversal mutations to the equivalent residues in mouse Otop1 (K149E, E219K in N domain, E433R, R586E in C domain) and tested their proton channel activity (Fig. 4k, l). The K149E mutant functioned similar to wildtype, while currents were greatly reduced in the E219K mutant, and not ‘rescued’ in the K149E/E219K double mutant. These results indicate that while introduction of a basic group at position 219 of mouse Otop1 is enough to disturb its functionality, the salt bridge interaction observed in the N domain vestibule of Otop1 is not critical for proper function. That the equivalent residues in Otop3 (Q146, E216) do not interact (Fig. 4e) further supports the notion that the apparent interaction in Otop1 is dispensable. Mutations in the C domain interaction yielded a different result; the greatly diminished current amplitude observed from each of the single mutants (E433R, R586E) was partly rescued in the double mutant (E433R/R586E). Therefore, this electrostatic interaction in the C-domain of Otop1 apparently supports proper proton channel function and the exact positioning of the residues is not essential. Indeed, the equivalent residues in Otop3 (Q435, R547) interact with each other in a similar fashion via apparent hydrogen bonding (Fig. 4j).

In addition to the salt bridges mentioned above, the N and C domains of Otop1 and Otop3 each contain a highly conserved glutamine-asparagine-tyrosine (QNY) triad (Q174/N204/Y268 and Q433/N528/Y571 in Otop1, Q175/N205/Y266 and Q429/N503/Y546 in Otop3) (Fig. 4d, e, i, j, Supplementary Fig. 1). Superposition of the domains shows that these triads occupy analogous positions (Supplementary Fig. 9a,b), and their side chains are close enough to interact directly or through intervening waters (Supplementary Fig. 9c-f).

### Molecular dynamics simulations show water penetration of Otop1

We conducted all-atom molecular dynamics (MD) simulations of Otop1 in a mixed lipid bilayer (80:20 POPC:CHOL) to examine areas susceptible to water penetration and thus to explore potential proton permeation pathways. As expected, the N and C domain vestibules both allowed water entry from the extracellular milieu into the membrane plane (Fig. 5a, b, d, e). Water flow through the central tunnel is completely blocked by the cholesterol molecules (Fig. 5c), supporting the notion that the tunnel lipids function as blockers of an extraneous permeation pathway. We observe the stochastic formation and breaking of a water wire in both the N and C domains (Supplementary Fig. 10). The breaking of the water wires occurs in the cytoplasmic half of the pathways, below the QNY triad, in part due to the presence of hydrophobic residues. Various hydrophilic side chains, including that of the QNY triad, contact water molecules and facilitate water penetration (Fig. 5d, e). The formation of a water wire during MD simulation suggests that proton conduction could be facilitated through a water-hopping mechanism. An average (two subunits and three simulations) distribution of water molecules along the putative pores (Fig. 5g) shows a marked difference in the degree of wetting between the extracellular and cytoplasmic halves of the pathways. An extended dry(er) region in the cytoplasmic half of the putative pores would not be expected to be conducive for proton conduction in the experimentally observed conformation, suggesting this might correspond to a hydrophobic gate region. However, we also note that proton conduction may not require a fully intact water wire^30^.

Surprisingly, the intrasubunit interface flanked by the N and C domains also permitted penetration of water from both sides of the bilayer, resulting in formation of a potential water wire. Most of the residues surrounding the water molecules at this interface are hydrophobic, with the notable exceptions of an arginine (R540), a glutamate (E267) and a histidine (H574) (Fig. 5f). E267 and H574 are highly conserved across Otopetrins (Supplementary Fig. 1). Though the cholesterol-like molecule present at the interface (Fig. 3c) is in close proximity to water molecules, its position in the Otop1 structure suggests it would only partially occlude the water wire (Fig. 5c). As noted above, the interactions of the cholesterol molecules in this region here are quite dynamic in our simulations, and the lipid molecules are able to diffuse away. In one simulation the hydrophobic tail of the cholesterol inserted further into the interface, causing a transient (∼5 ns) break in the water wire (Supplementary Fig. 10). On average the water distribution along the interface is more uniform than that of the N and C domains, without the clear break/hydrophobic gate seen in the latter. Overall, MD simulation suggests that the intrasubunit interface presents a third possible avenue for proton conduction in Otop1. That this interface is the most conserved region in Otopetrins (Supplementary Fig. 7a) further underscores its likely functional importance.

## Discussion

Despite their broad roles in biology, knowledge of proton channels has lagged behind that of other types of ion channels. The recent characterization of Otopetrins, previously implicated in vestibular function^11,12^, provides impetus for investigations of this novel proton channel family^10^. Here we have described cryo-EM structures of Otop1 and Otop3 in lipidic nanodiscs, elucidating the topology and dimeric organization of Otopetrins in a native-like bilayer environment. Our structures and MD simulations point to three possible routes for proton conduction per subunit: structurally similar openings in the N and C domains and the intrasubunit interface. At this point it remains uncertain which of these three pathways (or their combination) contributes to the proton currents recorded in electrophysiological experiments; we found mutations in both the N and C domain putative pores that result in loss of function. Clarifying the proton conduction pathway(s), as well as gating and selectivity mechanisms, will require further studies. Our results set the stage for such investigations on a family of proton channels involved in sour taste perception, vestibular function, and other as-yet identified roles.

## Acknowledgements

We thank W. Anderson for managing the electron microscopy facility at Scripps Research, H. Turner and J. Torres for help with data collection, C. Bowman for assistance with computation, D. Artiga and J. Kaplan for help with generating cDNA constructs and Vsevolod Katritch for providing expert advice on design of expression constructs. We acknowledge members of the Ward, Liman, Sansom, and Patapoutian labs for helpful advice. This work was supported by a Ray Thomas Edwards Foundation grant to A.B.W, funding from the NIH (NIDCD013741) to E.R.L, Wellcome (grant 208361/Z/17/Z), BBSRC (grants BB/N000145/1 and BB/R00126X/1), and EPSRC (grant EP/R004722/1) to M.S.P.S. K.S. is a postdoctoral fellow of the Jane Coffin Childs Memorial Fund for Medical Research. C.C.A.T. is supported by the Skaggs-Oxford Scholarship and the Croucher Foundation.

## Author contributions

K.S. prepared cryo-EM samples, collected and processed cryo-EM data, and built structures. B.T. performed electrophysiology experiments and analyzed data. C.C.A.T. performed molecular dynamics simulation and analyzed data. W.L. conducted FSEC experiments. Y.T. designed and generated constructs and analyzed data. M.S.P.S., E.R.L., and A.B.W. supervised molecular dynamics, functional experiments, and cryo-EM structure determination, respectively. K.S. drafted a majority of the manuscript, with significant additions from B.T., C.C.A.T., M.S.P.S., E.R.L., and A.B.W. All authors contributed to finalization of the manuscript.

## Data availability

Cryo-EM maps of zebrafish Otop1 and chicken Otop3 in nanodiscs have been deposited to the Electron Microscopy Data Bank under accession codes 9360 and 9361, respectively. Atomic coordinates of zebrafish Otop1 and chicken Otop3 have been deposited to the PDB under IDs 6NF4 and 6NF6, respectively. All other data are available upon reasonable request to the corresponding authors.

## Methods

### Constructs

Zebrafish Otop1 (zfOtop1, Uniprot entry Q7ZWK8) was obtained by PCR from zebrafish whole animal cDNA and confirmed by sequencing. Chicken Otop3 (chOtop3, Uniprot entry R4GK65) was synthesized and codon-optimized for expression in human cell lines. Both cDNAs were cloned into a pcDNA3.1 vector with an N-terminal fusion tag consisting of an octahistidine tag followed by eGFP, a Gly-Thr-Gly-Thr linker, and 3C protease cleavage site (LEVLFQGP). We call these constructs GFP-zfOtop1 and GFP-chOtop3 below. We note that the Uniprot entry for chicken Otop3 used in our study is a 561 amino acid sequence that has been updated since we had it synthesized to a 555 amino acid sequence. The two sequences are identical excepting a short region starting at residue 176. In our chOtop3 the sequence is ^176^AFFLWHHSKDCIQVQHNLTR^195^. The corresponding sequence in the updated Uniprot entry is ^176^VMQIPSWQLTHSL^189^. In addition, alanine at position 438 in the Uniprot entry is lysine in our sequence of chicken Otop3. The sequence we used is more highly conserved and likely to be correct. Mutations were introduced into the mouse Otop1 cDNA and sequenced as described in Tu et al^10^.

### FSEC and purification screening

Approximately one dozen Otopetrin orthologues fused to GFP at the N-terminus were screened by FSEC as previously described^19^. We identified GFP-zfOtop1 and GFP-chOtop3 as promising candidates for structure determination. To conduct FSEC, HEK293F cells suspended in Freestyle 293 expression medium were transfected with N-terminal GFP fusion constructs when they reached a cell density of ∼1.5 × 10^6^ / mL and incubated in an orbital shaker at 37° C supplemented with 8% CO_2_ for two days. 1 mL of cells was pelleted and resuspended in 200 µL of detergent-containing buffer and rotated at 4 °C for one hour, then centrifuged at greater than 50,000 × g in a tabletop ultracentrifuge, then injected onto a Superose 6 increase column (equilibrated to 20 mM Tris pH 8.0, 150 mM NaCl, 0.5 mM n-dodecyl-β-D-maltoside, 1 mM EDTA) in line with a fluorimeter tracking GFP fluorescence.

### Sample preparation

Similar procedures were used to prepare cryo-EM samples for zfOtop1 and chOtop3 in nanodiscs. HEK293F cells suspended in Freestyle 293 expression medium at a cell density of ∼1.5 × 10^6^ / mL were transfected with 1 mg of zfOTOP1 or chOTOP3 DNA and 3 mg polyethylenimines and incubated at 37°C for supplemented with 8% CO_2_ for 48 hours. Crude cell pellets were washed with cold PBS then resuspended in buffer containing 20 mM Tris pH 8.0, 150 mM NaCl, 1% n-dodecyl-β-D-maltoside, 0.15% cholesteryl hemisuccinate, 2 µg/µL leupeptin, 2 µg/µL aprotinin, 1 mM phenylmethylsulfonyl fluoride, 2 µM pepstatin A, 2 mM dithiothreitol (DTT) then stirred at 4 °C for 1 hour. The lysate was clarified by centrifugation in a JLA 16.25 rotor at 34,000 × *g* for 1 hour. Anti-GFP nanobody-coupled CNBr-Activated Sepharose 4B resin^31^ was added to the clarified lysate, and the mixture was rotated at 4°C for 1.5 hours. Resin was washed with more than 10 resin volumes of wash buffer (20 mM Tris pH 8.0, 150 mM NaCl, 0.07% n-dodecyl-β-D-maltoside, 0.01% cholesteryl hemisuccinate, 0.4 µg/µL aprotinin, 0.4 µM pepstatin A, 2 mM DTT). Washed resin was resuspended in wash buffer to make a ∼40% slurry and 50 ug of prescission protease was added, then incubated at 4°C for five hours. Flowthrough containing cleaved zfOtop1 or chOtop3 was collected, concentrated in 100 kDa cutoff centrifugal filter, then injected into a Superose 6 increase column equilibrated with wash buffer. Peak fractions corresponding to zfOtop1 or chOtop3 were collected and concentrated to ∼1 mg/mL, then mixed with MSP2N2 nanodisc scaffold protein^32^ and soybean polar lipid extract at a molar ratio of approximately 1:1.67:13.33 (monomer:MSP2N2:lipids) for zfOTOP1 and approximately 1:2.67:26.67 for chOTOP3. The total volume of the mixture was approximately 0.5 mL. The mixture was incubated on ice for one hour, then 100 mg of biobeads was added and nanodisc assembly was initiated by rotation overnight at 4° C. After removing biobeads by centrifugation, the nanodisc assembly mixture was injected into a Superose 6 increase column equilibrated with buffer containing 20 mM Tris pH 8.0, 150 mM NaCl, 0.4 µg/µL aprotinin, 0.4 µM pepstatin A, 2 mM DTT. Peak fractions were concentrated to 2.1 mg/mL (for zfOtop1) or 1.1 mg/mL (for chOtop3) using a 100 kDa cutoff centrifugation filter. 3.5 μL of purified protein was applied to previously plasma-cleaned UltrAuFoil 1.2/1.3 300 mesh grids and blotted once for 3.5 seconds with blot force 0 after a wait time of 10 seconds. Blotted grids were plunge frozen into nitrogen-cooled liquid ethane using a Vitrobot Mark IV operated at 10 °C and 100% humidity.

### Cryo-EM data collection

Images were collected at 300 kV using a Titan Krios coupled with a K2 Summit direct electron detector (Gatan) at a nominal magnification of 29,000x with a pixel size of 1.03 Å. 53 frames (for zfOtop1) or 45 frames (for chOTOP3) of 250 ms exposure time were collected per movie, resulting in a total accumulated dose of ∼50 electrons per Å^2^. Automated micrograph collection was performed using Leginon software^33^ with a target defocus range of 0.6-1.8 µM.

### Cryo-EM Data Processing

Micrographs were aligned and dose-weighted using MotionCor2^34^. Contrast transfer function (CTF) parameters were obtained with Gctf^35^. Data processing for zfOtop1 and chOtop3 were carried out similarly. The following processing procedure was used for zfOtop1; the procedure for chOtop3 is depicted as a flow chart in Supplementary Fig. 4d. 1,517,125 particles were picked from 2,138 micrographs using AutoPick in RELION-2.1^36^ applying a gaussian blob as a reference. Picked particles were extracted without binning using RELION-2.1 with box size of 200 pixels (206 Å) and exported to cryoSPARC v0.6.5^37^. Contaminants and particles not containing features of zfOTOP1 were removed by two rounds of 2D classification, resulting in 792,311 particle images that were retained for further processing. These particles were then subjected to *ab initio* reconstruction without symmetry, requesting 3 classes and a maximum resolution of 7 Å. The most populated class contained 435,468 particles and had secondary structure features apparent. These particles were transferred to RELION-2.1 and refined with C2 symmetry applied and using the *ab initio* reconstruction low pass filtered to 12 Å resolution as the initial reference. This refinement resulted in a 3.26 Å resolution map after postprocessing that displayed anisotropic features. The particles were then recentered and reextracted on the basis of their refined coordinates, followed by 3D classification without alignment using *k*=6 and tau fudge value of 8. This classification run was repeated twice, and the particles from the highest resolution class from each run were combined and duplicates removed, resulting in 76,546 particles. Masked refinement of these particles resulted in a 3.16 Å resolution map after postprocessing that did not exhibit anisotropic features. The refined particles were then subjected to local CTF and beam tilt estimation using the ctfrefine jobtype in RELION-3.0^38^. Subsequent masked refinement of these particles resulted in a 3.07 Å resolution map. The refinement parameters and reconstruction from this refinement run were used as inputs for Bayesian polishing in RELION-3.0^38^. After bayesian polishing, masked refinement yielded a 3.02 Å resolution reconstruction. The polished particles were then subjected to a final round of 3D classification without alignment (k=3, tau fudge=8). Masked refinement of 67,425 particles from the highest resolution class resulted in the final 2.98 Å resolution map. For all refinements and classification jobs, C2 symmetry was applied unless otherwise noted, and stated resolutions were calculated at FSC=0.143 by postprocessing in RELION using a soft mask around the zfOtop1 protein and nanodisc belt.

### Model building, refinement, and visualization

Cryo-EM maps were flipped to the correct handedness using the ‘vop zflip’ command in UCSF chimera^39^ and sharpened with phenix.autosharpen^40^ prior to model building in coot^41^. Manual building was iterated with real space refinement using phenix^42^ and rosetta^43^. The zfOtop1 model was built first. Most side chains in the TM helices were clearly resolved, allowing *de novo* model building using bioinformatic prediction of the positions of TM segments^44^ as a guide. To build the chOtop3 model, a homology model of chOtop3 was produced from the zfOtop1 structure using SWISS-MODEL^45^ and docked into the Otop3 map, then manually adjusted and refined. Geometric restraints for cholesterol and CHS molecules were produced using the ELBOW program in phenix. The distal N-terminus and portions of linkers connecting TM2-TM3, TM5-TM6, TM6-TM7, TM10-TM11 were omitted from either model due to poor density. The model of zfOtop1 (586 residues in full-length) contains residues 46-113, 123-221, 248-283, 302-443, 510-585. The model of Otop3 (561 residues in full-length) contains residues 49-116, 124-227, 246-283, 304-366, 375-438, 487-560. Structural figures were made in PyMOL^46^, UCSF Chimera^47^, or UCSF ChimeraX^48^. Sequence conservation scores were calculated and mapped onto the structure of zfOtop1 using Consurf^49,50^, with ∼100 Otopetrin sequences obtained from UniprotKB as input, aiming for even distribution between Otop1, Otop2, and Otop3 subtypes. Sequence alignment was calculated using Clustal Omega^51^ and represented using ESPript 3.0^52^.

### Cell culture and transfection for electrophysiology

HEK-293 cells (CRL-1573, ATCC) were cultured in DMEM containing 10% fetal calf serum and 50 μg/ml gentamycin. Cells were transfected in 35mm petri dishes, with approximately 600 ng DNA and 2 μl TransIT-LT1 transfection reagent (Mirus Bio Corporation) following manufacturer’s protocol. Constructs were expressed as N-terminal GFP fusion (for zfOtop1 and chOtop3, see above) or a N-terminal YFP fusion (mOtop1 mutants). Fluorescence was used to select transfected cells. The cells were lifted using Trypsin-EDTA 24h after transfection and plated onto a coverslip for patch clamp recordings.

### Patch clamp electrophysiology

Whole-cell patch clamp recording was performed as previously described ^10^. Briefly, recordings were made with an Axonpatch 200B or 700B amplifier, digitized with a Digidata 1322a 16-bit data acquisition system, acquired with pClamp 8.2 and analyzed with Clampfit 8.2 (Molecular devices, Palo Alto, CA). Records were sampled at 5 kHz and filtered at 1 kHz. Patch pipettes with resistance of 2 – 4 MΩ were fabricated from borosilicate glass and only recordings in which a gigaohm seal was achieved were used in the analysis. For most of the experiments, the membrane potential was held at −80 mV or ramped from −80 mV to +80 mV (1V/s) once per second and solutions were exchanged with a Warner fast step system.

### Patch clamp electrophysiology solutions

Standard pipette solution contained 120 mM Cs-aspartate, 15 mM CsCl, 2 mM Mg-ATP, 5 mM EGTA, 2.4 mM CaCl2 (100 nM free Ca2+), and 10 mM HEPES (pH 7.3 with CsOH; 290 mosm). Proton currents were evoked in Na^+^-free, NMDG^+^-based extracellular solutions that contained 160 mM NMDG-Cl, 2mM CaCl2, and either 10 mM HEPES (for pH 7.4), 10 mM MES (for pH 6-5.5), or 10 mM HomoPIPES (for pH 5-4.5), pH adjusted with HCl. ZnCl_2_ was added to these solutions, and pH adjusted at concentrations indicated. Prior to recording, cells were bathed in Tyrode’s solution contained 145 mM NaCl, 5 mM KCl, 1 mM MgCl2, 2 mM CaCl2, 20mM D-Glucose, 10mM HEPES (pH 7.4 with NaOH).

### Building the atomistic MD simulation system

All molecular dynamics simulations were carried out using GROMACS 5.0.2^53^ on the nanodisc-embedded zfOtop1 structure. The cholesterol molecules associated with the structure were first removed, then converted into a coarse-grained (CG) representation (force field: MARTINI 2.2)^54^ using MemProtMD^55^. The protein was embedded in a band of either 100% POPC or 80%/20% POPC/cholesterol molecules that are randomly oriented, then the periodic simulation box was completed with the addition of water and 0.15 M NaCl. A 100-ns CG simulation with protein backbone beads position restrained (force constant: 1000 kJ mol^−1^ nm^−2^) permitted lipid self-assembly. The final frame was then converted to atomistic (AT) detail using CG2AT-Align^56^, with the CHARMM36 force field^57^ and solvated again in TIP3P water^58^ and 0.15 M NaCl. For the set of simulations in the mixed membrane, the eight cholesterol molecules associated with zfOtop1 were manually added back to the AT system, with water and lipid molecules that resulted in steric clashes removed.

### Atomistic MD simulations

The system with added the cholesterol molecules was energy minimised to maximum force 1000 kJ mol^−1^ nm^−2^, before running a 1-ns simulation (time-step: 1 fs) with the protein backbone atoms position restrained (force constant: 1000 kJ mol^−1^ nm^−2^). Three repeats of 100-ns production run (time-step: 2 fs) were then performed based on the final snapshot of the short simulation, without position restraints. Here, distance restraints were applied between endings of the missing loops (force constant: 1000 kJ mol^−1^ nm^−2^; lower and upper bound values *r*_0_ and *r*_1_ were ±1 Å of the starting distances). The AT simulations were performed as NPT ensembles held at 1 bar and 310 K. A semi-isotropic Parrinello-Rahman barostat^59^ (coupling constant: 1 ps; compressibility: 4.5×10%^−5^ bar^−1^) and a velocity-rescaling thermostat^60^ (coupling constant: 0.1 ps) were used. All covalent bonds were constrained using the LINCS algorithm^61^. Electrostatics were modelled with a Particle Mesh Ewald (PME) model^62^, and van der Waals’ interactions were modelled using a cut-off scheme, both with cut-off distances at 10 Å.

### MD simulations trajectory analysis

Root-mean-square deviations measurements were performed using the *RMSD Trajectory Tool* in VMD^63^. Protein-cholesterol distances were calculated with the *distance* tool in GROMACS based on their centres of geometry. For the extraction of water trajectories, water molecules within the N domain pathway were selected based on their mutual proximity to TM3, TM4 and TM6, while excluding those found in bulk water. A cuboid selection (13×18.5×42 Å^3^) was used to select water molecules within the C domain pathway. For the intrasubunit interface, a similar cuboid method (10×12×42 Å^3^) was used and it covers the outer half of interface. MD-related figures were rendered using PyMOL^46^.

