## Supplementary Information for "Structures of the Otopetrin Proton Channels Otop1 and Otop3"

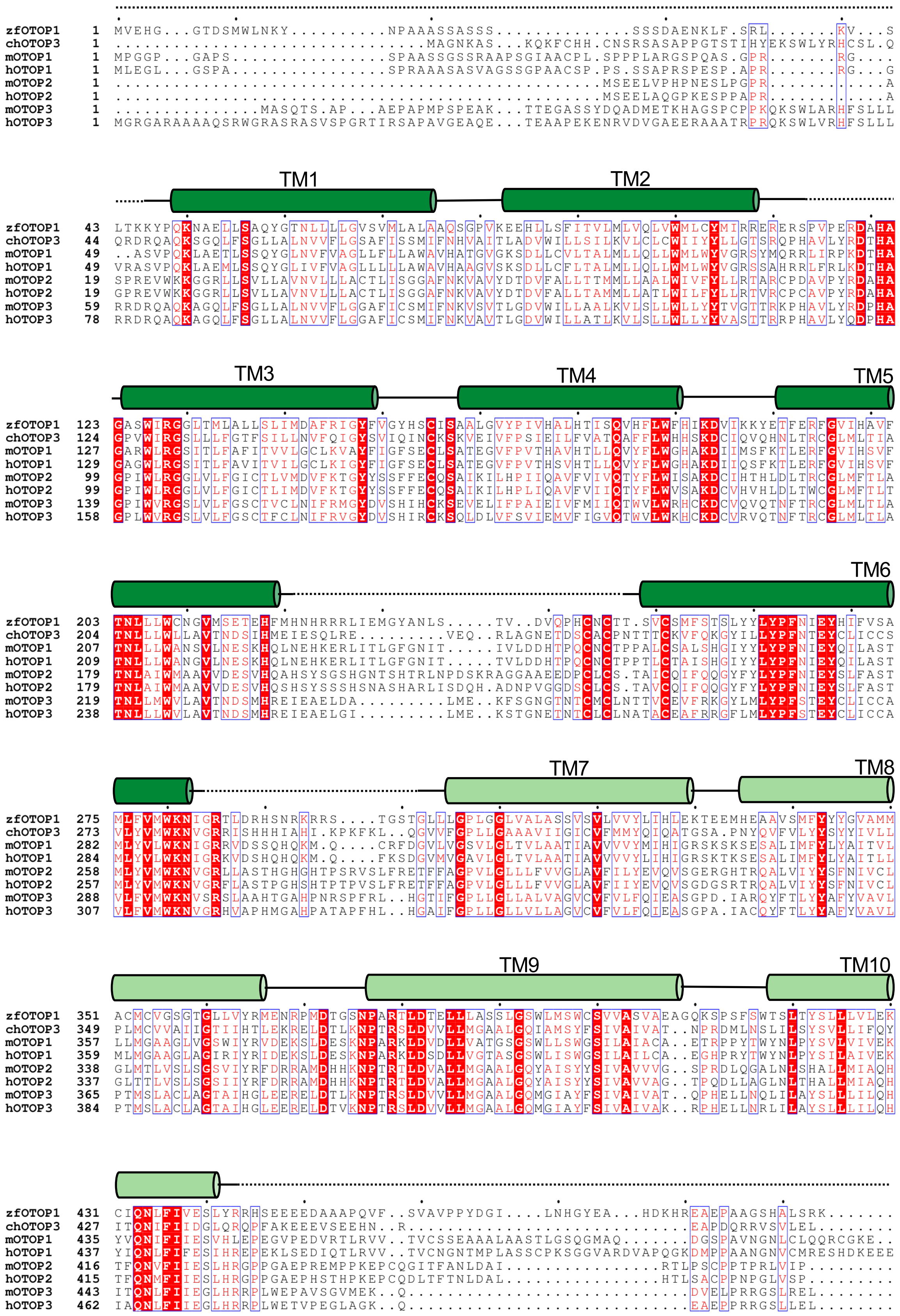
zfOTOP1 1 MVEHG...GTDSMWLNKY.........NPAAASSASSS.............SSSDAENKLF.SRL.......KV...S
chOTOP3 1 ...........................MAGNKAS...KQKFCHH.CNSRSASAPPGTSTIHYEKSWLYRHCSL.Q
mOTOP1 1 MPGGP...GAPS...............SPAASSGSSRAAPSGIAACPL.SPPPLARGSPQAS.GPR.......RG....
hOTOP1 1 MLEGL...GSPA...............SPRAAASASVAGSSGPAACSP.PS.SSAPRSPESP.APR.......RG...G
mOTOP2 1 ..............................................MSEELVPHPNESLPGPR...........A
hOTOP2 1 ..............................................MSEELAQGPKESPPAPR...........A
mOTOP3 1 ..............MASQTS.AP...AEPAPMPSPEAK...TTEGASSYDQADMETKHAGSPCPPKQKSWLARHFSLLL
hOTOP3 1 MGRGARAAAAQSRWGRASRASVSPGRTIRSAPAVGEAQE...TEAAPEKENRVDVGAEERAAATRPRQKSWLVRHFSLLL
TM1
TM2
zfOTOP1 43 LTKKYPQKNAELLSAQYGTNLLLLGVSVMLALAAQSGPVKEEHLLSFITVLMLVQLVWMLCYMIRRERERSPVPERDAHA
chOTOP3 44 QRDRQAQKSGQLFSGLLALNVVFLGSAFISSMIFNHVAITLADVWILLSILKVLCLCWIIYYLLGTSRQPHAVLYRDTHA
mOTOP1 49 ..ASVPQKLAETLSSQYGLNVFVAGLLFLLAWAVHATGVGKSDLLCVLTALMLLQLLWMLWYVGRSYMQRRLIRPKDTHA
hOTOP1 49 VRASVPQKLAEMLSSQYGLIVFVAGLLLLLAWAVHAAGVSKSDLLCFLTALMLLQLLWMLWYVGRSSAHRRLFRLKDTHA
mOTOP2 19 SPREVWKKGGRLLSVLLAVNVLLLACTLISGGAFNKVAVYDTDVFALLTTMMLLAALWIVFYLLRTARCPDAVPYRDAHA
hOTOP2 19 GPREVWKKGGRLLSVLLAVNVLLLACTLISGGAFNKVAVYDTDVFALLTAMMLLATLWILFYLLRTVRCPCAVPYRDAHA
mOTOP3 59 RRDRQAQKAGQLFSGLLALNVVFLGGAFICSMIFNKVSVTLGDVWILLAALKVLSLLWLLYYTVGTTRKPHAVLYRDPHA
hOTOP3 78 RRDRQAQKAGQLFSGLLALNVVFLGGAFICSMIFNKVAVTLGDVWILLATLKVLSLLWLLYYVASTTRRPHAVLYQDPHA
TM3
TM4
TM5
zfOTOP1 123 GASWIRGGLTMLALLSLIMDAFRIGYFVGYHSCISAALGVYPIVHALHTISQVHFLWFHIKDVIKKYETFERFGVIHAVF
chOTOP3 124 GPVWIRGSLLLFGTFSILLNVFQIGYSVIQINCKSKVEIVFPSIEILFVATQAFFLWHHSKDCIQVQHNLTRCGLMLTIA
mOTOP1 127 GARWLRGSITLFAFITVVLGCLKVAYFIGFSECLSATEGVFPVTHAVHTLLQVYFLWGHAKDIIMSFKTLERFGVIHSVF
hOTOP1 129 GAGWLRGSITLFAVITVILGCLKIGYFIGFSECLSATEGVFPVTHSVHTLLQVYFLWGHAKDIIQSFKTLERFGVIHSVF
mOTOP2 99 GPIWLRGGLVLFGICTLVMDVFKTGYYSSFFECQSAIKILHPIIQAVFVIVQTYFLWISAKDCIHTHLDLTRCGLMFTLA
hOTOP2 99 GPIWLRGGLVLFGICTLIMDVFKTGYYSSFFECQSAIKILHPLIQAVFVIIQTYFLWVSAKDCVHVHLDLTWCGLMFTLT
mOTOP3 139 GPIWVRGSLVLFGSCTVCLNIFRMGYDVSHIHCKSEVELIFPAIEIVFMIIQTWVLWRHCKDCVQVQTNFTRCGLMLTLA
hOTOP3 158 GPLWVRGSLVLFGSCTFCLNIFRVGYDVSHIRCKSQLDLVFSVIEMVFIGVQTWVLWKHCKDCVRVQTNFTRCGLMLTLA
TM6
zfOTOP1 203 TNLLLWCNGVMSETEHFMHNHRRRLIEMGYANLS.....TV..DVQPHCNCTT.SVCSMFSTSLYYLYPFNIEYHIFVSA
chOTOP3 204 TNLLLWLLAVTNDSIHMEIESQLRE.........VEQ..RLAGNETDSCACPNTTTCKVFQKGYILLYPFNTEYCLICCS
mOTOP1 207 TNLLWANSVLNESKHQLNEHKERLITLGFGNIT.....IVLDDHTPQCNCTPPALCSALSHGIYYLYPFNIEYQILAST
hOTOP1 209 TNLLWANGVLNESKHQLNEHKERLITLGFGNIT.....TVLDDHTPQCNCTPPTLCTAISHGIYYLYPFNIEYQILAST
mOTOP2 179 TNLAIWMAAVVDESVHQAHSYSGSHGNTSHTRLNPDSKRAGGAAEEDPCLCS.TAICQIFQQGYFYLYPFNIEYSLFAST
hOTOP2 179 TNLAIWMAAVVDESVHQSHSYSSSHSNASHARLISDQH.ADNPVGGDSCLCS.TAVCQIFQQGYFYLYPFNIEYSLFAST
mOTOP3 219 TNLLMWVLAVTNDSMHREIEAELDA.........LME..KFSGNGTNTCMCLNTTVCEVFRKGYLMLYPFSTEYCLICCA
hOTOP3 238 TNLLLWVLAVTNDSMHREIEAELGI.........LME..KSTGNETNTCLCLNATACEAFRRGFLMLYPFSTEYCLICCA
TM7
TM8
zfOTOP1 275 MLFVMWKNIGRTLDRHSNRKRRS....TGSTGLLLGPLGGLVALASSVSVLVVYLIHLEKTEEMHEAAVSMFYYYGVAMM
chOTOP3 273 VLYVMWKNVGRRISHHHIAHI.KPKFKL..QGVVFGPLLGAAAVIIGICVFMMYQIQATGSA.PNYQVFVLYYSYYIVLL
mOTOP1 282 MLYVLWKNIGRRVDSSQHQKM.Q....CRFDGVLVGSVLGLTVLAATIAVVVVYMIHIGRSKSKSESALIMFYLYAITVL
hOTOP1 284 MLYVLWKNIGRKVDSHQHQKM.Q....FKSDGVMVGAVLGLTVLAATIAVVVVYLIHIGRSKTKSESALIMFYLYAITLL
mOTOP2 258 MLYVMWKNVGRLLASTHGHGHTPSRVSLFRETFFAGPVLGLLLFVVGLAVFILYEVQVSGERGHTRQALVIYYSFNIVCL
hOTOP2 257 MLYVMWKNVGRFLASTPGHSHTPTPVSLFRETFFAGPVLGLLLFVVGLAVFIIYEVQVSGDGSRTRQALVIYYSFNIVCL
mOTOP3 288 VLFVMWKNVSRSLAAHTGAHPNRSPFRL..HGTIFGPLLGLLALVAGVCVFVLFQIEASGPD.IARQYFTLYYAFYVAVL
hOTOP3 307 VLFVMWKNVGRHVAPHMGAHPATAPFHL..HGAIFGPLLGLLVLLAGVCVFVLFQIEASGPA.IACQYFTLYYAFYVAVL
TM9
TM10
zfOTOP1 351 ACMCVGSGTGLLVYRMENRPMDTGSNPARTLDTELLLASSLGSWLMSWCSVVASVAEAGQKSPSFSWTSLTYSLLLVLEK
chOTOP3 349 PLMCVVAIIGTIIHTLEKRELDTLKNPTRSLDVVLLMGAALGQIAMSYFSIVAIVAT..NPRDMLNSLILSYSVLLIFQY
mOTOP1 357 LLMGAAGLVGSWIYRVDEKSLDESKNPARKLDVDLLVATGSGSWLLSWGSILAIACA..ETRPPYTWYNLPYSVLVIVEK
hOTOP1 359 MLMGAAGLAGIRIYRIDEKSLDESKNPARKLDSDLLVGTASGSWLISWGSILAILCA..EGHPRYTWYNLPYSILAIVEK
mOTOP2 338 GLMTLVSLSGSVIYRFDRRAMDHHKNPTRTLDVALLMGAALGQYAISYYSIVAVVVG..SPRDLQGALNLSHALLMIAQH
hOTOP2 337 GLTTLVSLSGSIIYRFDRRAMDHHKNPTRTLDVALLMGAALGQYAISYYSIVAVVAG..TPQDLLAGLNLTHALLMIAQH
mOTOP3 365 PTMSLACLAGTAIHGLEERELDTLKNPTRSLDVVLLMGAALGQMGIAYFSIVAIVAT..QPHELLNQLILAYSLLLILQH
hOTOP3 384 PTMSLACLAGTAIHGLEERELDTVKNPTRSLDVVLLMGAALGQMGIAYFSIVAIVAK..RPHELLNRLILAYSLLLILQH
zfOTOP1 431 CIQNLFIVESLYRRHSEEEEDAAAPQVF..SVAVPPYDGI....LNHGYEA...HDKHREAEPAAGSHALSRK.......
chOTOP3 427 ITQNIFIIDGLQRQPFAKEEEVSEEHN..R...........................EAPDQRRVSVLEL........
mOTOP1 435 YVQNIFIIESVHLEPEGVPEDVRTLRVV..TVCSSEAAALAASTLGSGQGMAQ.......DGSPAVNGNLCLQQRCGKE..
hOTOP1 437 YIQNLFIFESIHREPEKLSEDIQTLRVV..TVCNGNTMPLASSCPKSGGVARDVAPQGKDMPPAANGNVCMRESHDKEEE
mOTOP2 416 TFQNVFIIESLHRGPPGAEPREMPPKEPCQGITFANLDAI.................RTLPSCPPTPRLVIP........
hOTOP2 415 TFQNMFIIESLHRGPPGAEPHSTHPKEPCQDLTFTNLDAL.................HTLSACPPNPGLVSP........
mOTOP3 443 ITQNLFIIEGLHRRPLWEPAVSGVMEK..Q..........................DVELPRRGSLREL........
hOTOP3 462 IAQNLFIIEGLHRRPLWETVPEGLAGK..Q..........................EAEPRRGSLLEL........

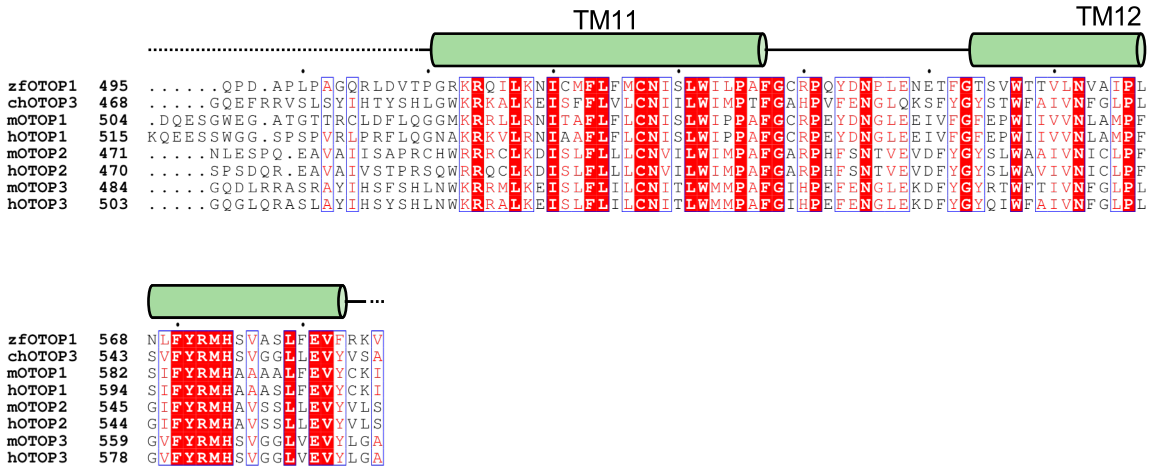


**Supplementary Figure 1. Topology and sequence alignment of Otopetrin subtypes.** zfOtop1 and chOtop3 were aligned with mouse Otop1 (mOTOP1, Uniprot ID Q80VM9), human Otop1 (hOTOP1, Uniprot ID Q7RTM1), mouse Otop2 (mOTOP2, Uniprot ID Q80SX5), human Otop2 (hOTOP2, Uniprot ID Q7RTS6), mouse Otop3 (mOTOP3, Uniprot ID Q80UF9), human Otop3 (hOTOP3, Uniprot ID Q7RTS5). The transmembrane helices in the zfOtop1 structure are depicted as cylinders above the sequence. Linkers that were modeled in the zfOtop1 structure are depicted as solid lines, while portions of the sequence excluded from the model due to poor density are depicted as dotted lines.

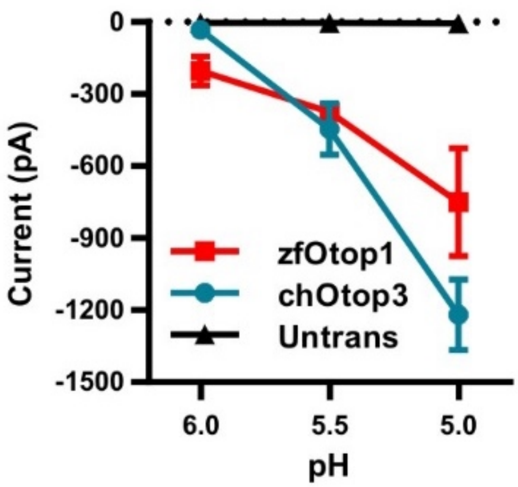


**Supplementary Figure 2. Proton channel properties of zfOtop1 and chOtop3.** Current magnitude as a function of pH in HEK-293 cells expressing zfOtop1 (red squares, n = 6), chOtop3 (cyan circle, n = 4) and in untransfected control cells (black triangles, n = 4). Error bars show s.e.m.

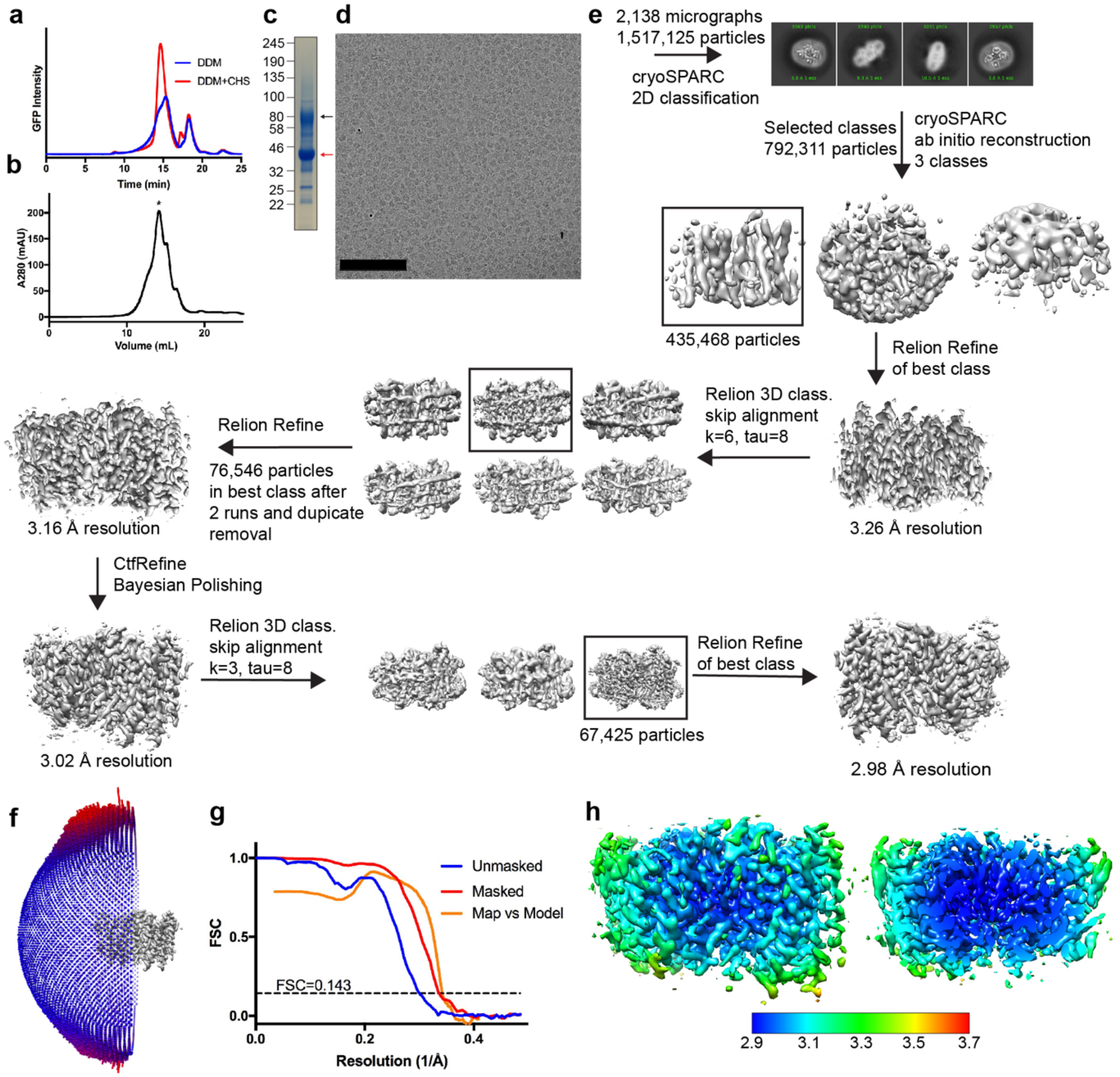


**Supplementary Figure 3. Cryo-EM data processing for Otop1. a,** FSEC traces of HEK cells expressing N-terminal GFP fusion of Otop1 solubilized in buffer containing 1% DDM (blue) or 1% DDM and 0.15% CHS (red). **b,** preparative SEC trace of Otop1 reconstituted in MSP2N2 nanodisc containing soybean lipids. **c,** SDS-PAGE analysis of purified Otop1 reconstituted in nanodiscs that was used for cryo-EM. Black arrow points to band corresponding to Otop1, while the red arrow points to band corresponding to MSP2N2 scaffold. **d,** representative aligned and dose-weighted cryo-EM micrograph. Scale bar = 100 nM. **e**, cryo-EM data processing scheme to obtain final reconstruction. **f**, relative angular distribution of final reconstruction used for model building. Red bars represent views with more particles. **g**, FSC plots of unmasked (blue) and masked (red) cryo-EM reconstructions, and model vs map comparison calculated in Phenix 1.14. (orange). **h,** side view (left) and centrally sliced side view (right) of Otop1 cryo-EM map sharpened to a b-factor of -80 Å$^2$ and colored according to local resolution determined by blocres program in RELION 3.0. Color key shows local resolution in Å.

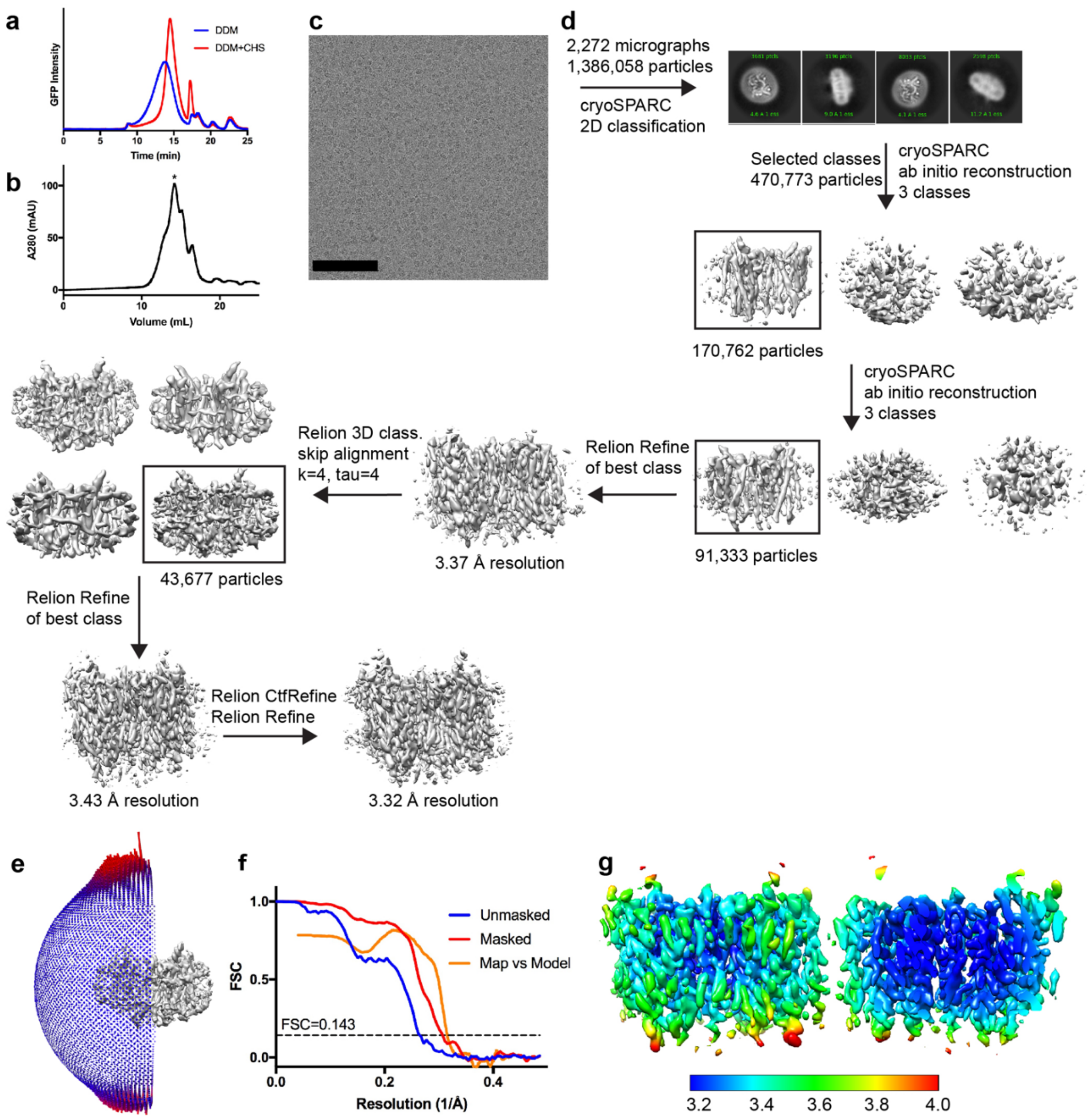


**Supplementary Figure 4. Cryo-EM data processing for Otop3. a,** FSEC traces of HEK cells expressing N-terminal GFP fusion of Otop3 solubilized in buffer containing 1% DDM (blue) or 1% DDM and 0.15% CHS (red). **b**, preparative SEC trace of Otop1 reconstituted in MSP2N2 nanodisc containing soybean lipids. **c**, representative aligned and dose-weighted cryo-EM micrograph. Scale bar = 100 nM. **d,** cryo-EM data processing scheme to obtain final reconstruction. **e**, relative angular distribution of final reconstruction used for model building. Red bars represent views with more particles. **f**, FSC plots of unmasked (blue) and masked (red) reconstructions, and model vs map comparison (orange). **g**, side view (left) and centrally sliced side view (right) of Otop3 cryo-EM map sharpened to a b-factor of -80 Å$^2$ and colored according to local resolution determined by blocres program in RELION 3.0. Color key shows local resolution in Å.

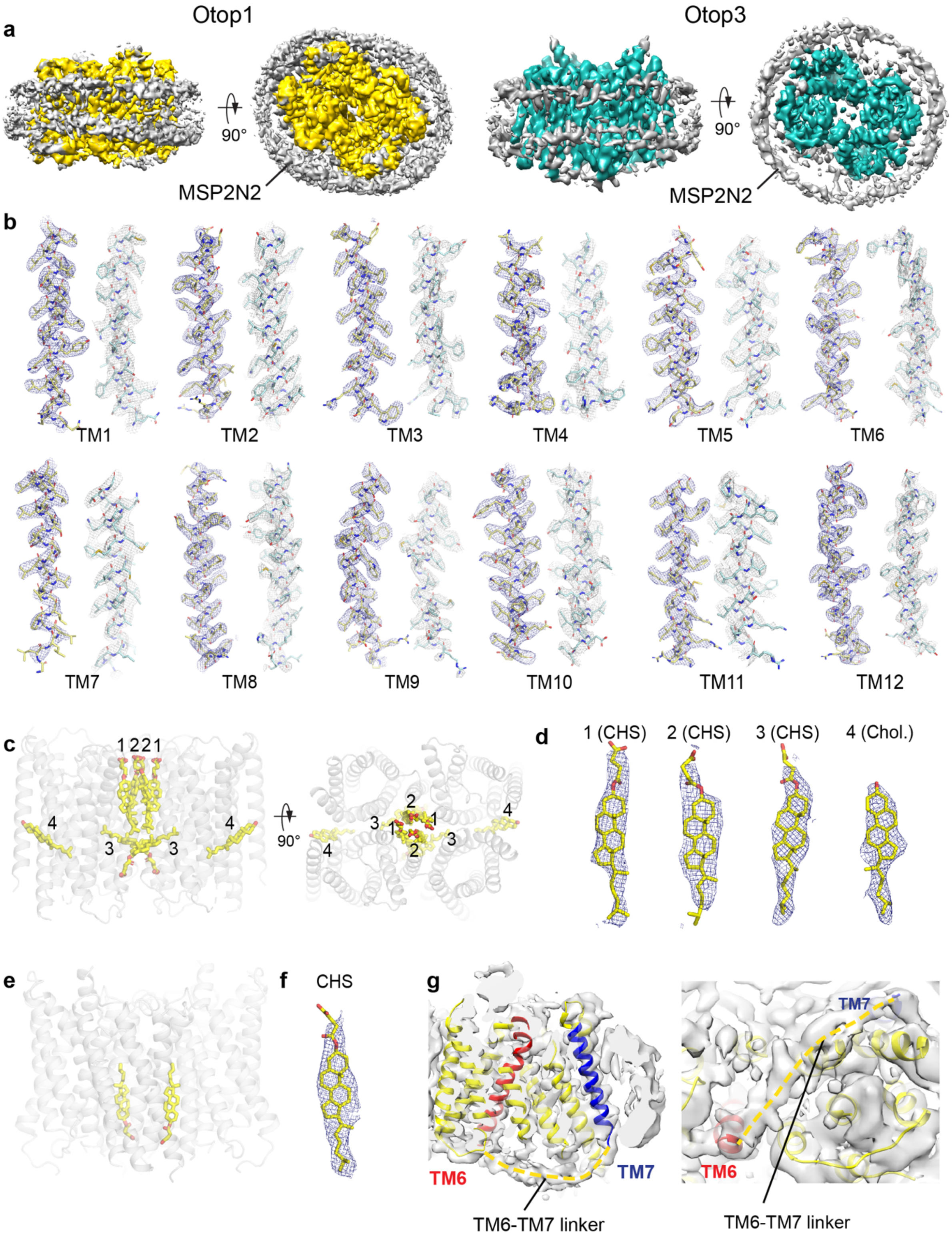


**Supplementary Figure 5. Fit of cryo-EM density to molecular model of Otop1 and Otop3.** **a**, side and top views of unsharpened maps of Otop1 (left) and Otop3 (right). Density within 2 Å of the molecular model is colored yellow (Otop1) or cyan (Otop3). Density for the MSP2N2 scaffold protein can be observed (gray density**). b,** isolated atomic model fragments of Otop1 (yellow) and Otop3 (cyan) with sharpened maps of Otop1 (blue mesh, 5 σ) or Otop3 (gray mesh,

4.5 σ) superimposed. **c**, Side (left) and top (right) views of molecular model of the Otop1 dimer, with bound CHS or cholesterol molecules depicted as sticks and numbered. **d**, sharpened cryo-EM maps of Otop1 superimposed onto CHS or cholesterols shown in C. σ values for the maps generated in Pymol are as follows: #1, 4 σ, #2 = 2.5 σ, #3 = 3 σ, #4 = 3 σ. **e**, side view of Otop3 dimer model with modeled CHS molecules shown as yellow sticks. **f**, Sharpened map of Otop3 at 3 σ is superimposed onto the CHS molecule. **g**, unsharpened cryo-EM map of Otop1 low-pass filtered to 4 Å resolution reveals weak density for the TM6-TM7 linker (highlighted by dashed yellow line), demonstrating the connectivity of the N and C domains within a single subunit. 18 residues are missing from the atomic model in the TM6-TM7 linker, which is sufficient to span this distance as an extended coil.

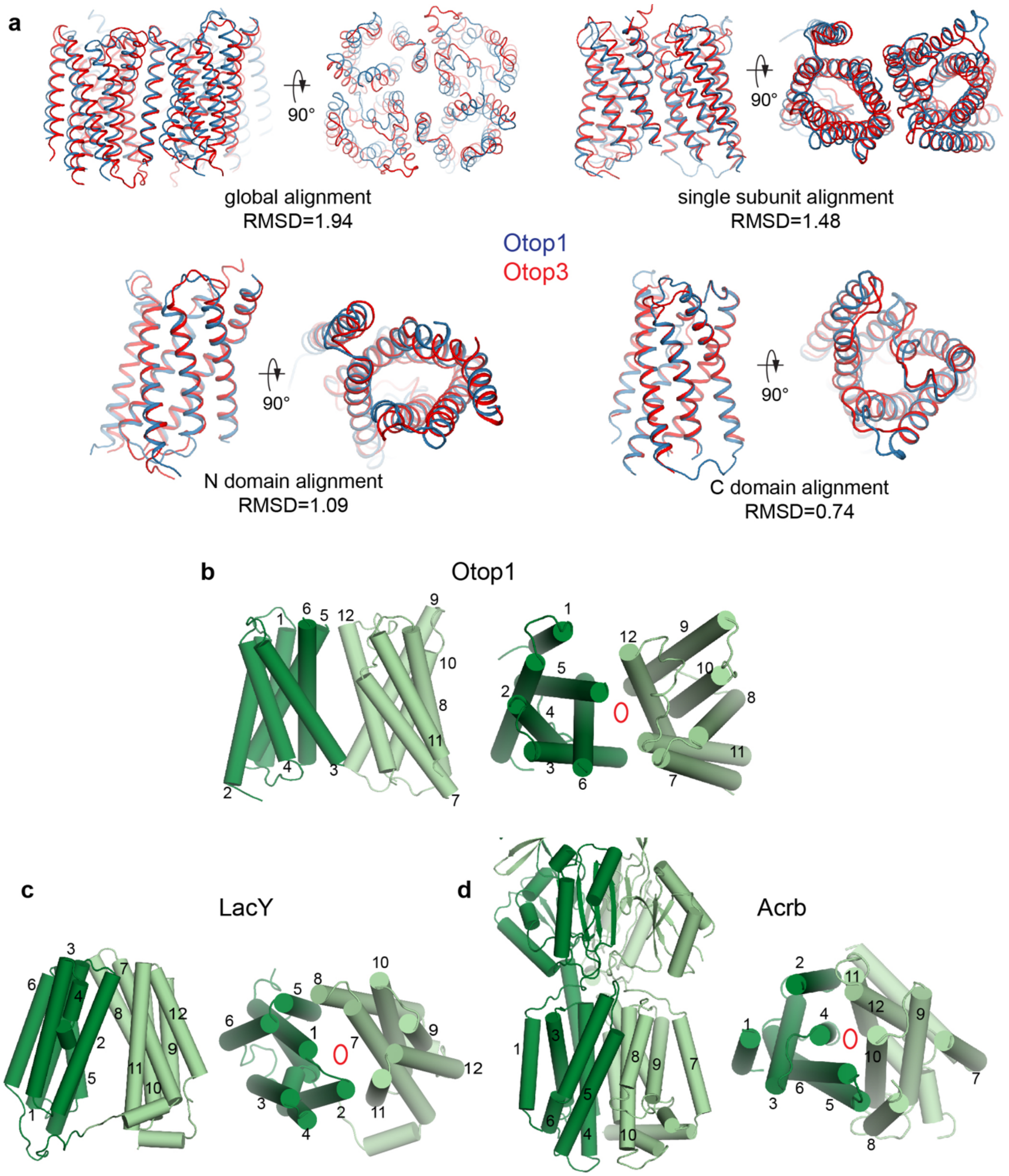


**Supplementary Figure 6. Structural comparison of Otop1 to Otop3, and Otop1 to LacY, a canonical MFS transporter, and Acrb, a canonical RND protein. a,** Structural alignments of Otop1 (blue) and Otop3 (red) based on the entire dimer, a single subunit, N domain, or C domain. Alignments and RMSD values were calculated using the 'align' program in Pymol. **b-d**, Side and top views of the Otop1 subunit (**b**), LacY lactose permease (**c**, PDB ID 1PV6), and Acrb multidrug efflux transporter (**d**, PDB ID 1IWG). Like Otopetrins, LacY and Acrb have twelve TM helices split into two 6-TM helix-containing bundles, the N (green) and C (light green) domains related by a two-fold pseudosymmetry axis (red ellipsoids). The TM helices are numbered to show that despite the similarity in domain organization, the folds of Otop1, LacY, and Acrb are distinct.

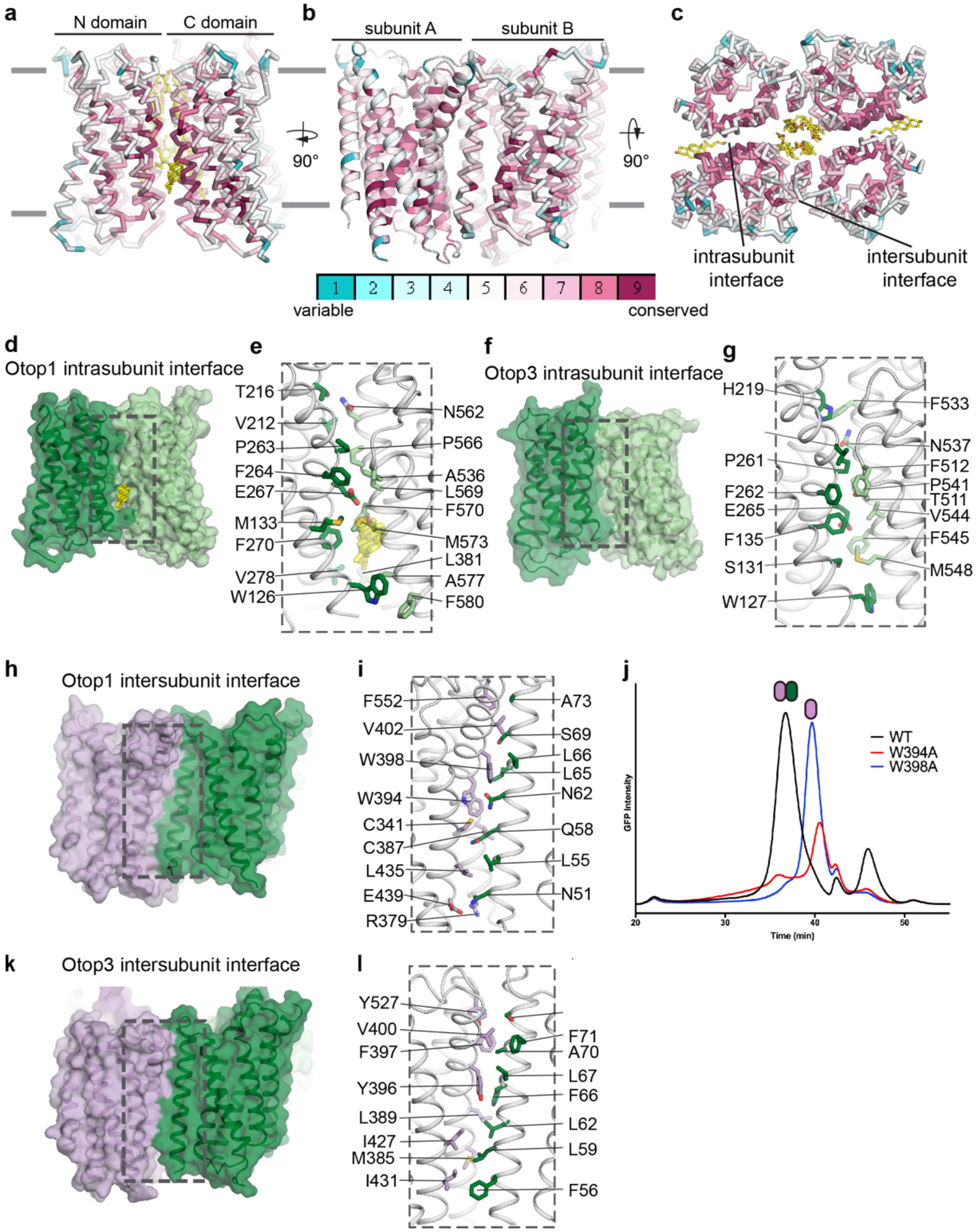


**Supplementary Figure 7. Intersubunit and intrasubunit interfaces. a-c,** side (**a,b**) and top (**c**) views of the Otop1 dimer, colored by Consurf sequence conservation score. Magenta represents more conserved, and teal represents less conserved. Bound cholesterol molecules are depicted as yellow sticks. **d-g**, representations of the intrasubunit interface in Otop1 (**d**) and Otop3 (**f**). **e,g** show expanded views of the interface with interfacial side chains shown in stick representation and labeled. The bound cholesterol molecule at the intrasubunit interface in Otop1 is shown. **h,** model of Otop1, with one subunit colored purple and the other subunit colored green. **i,** expanded view of boxed region in **h**, with residues contributing to intersubunit shown as stick. **j,** FSEC analysis of GFP-fused wild type zebrafish Otop1 and constructs harboring alanine mutations at two tryptophans (W394A, W398A) at the intersubunit interface. The mutants display right-shifted peaks compared to the wild type dimeric peak, consistent with solubilization of monomeric species. **k, l,** overall (**k**) and expanded (**l**) views of Otop3 intersubunit interface.

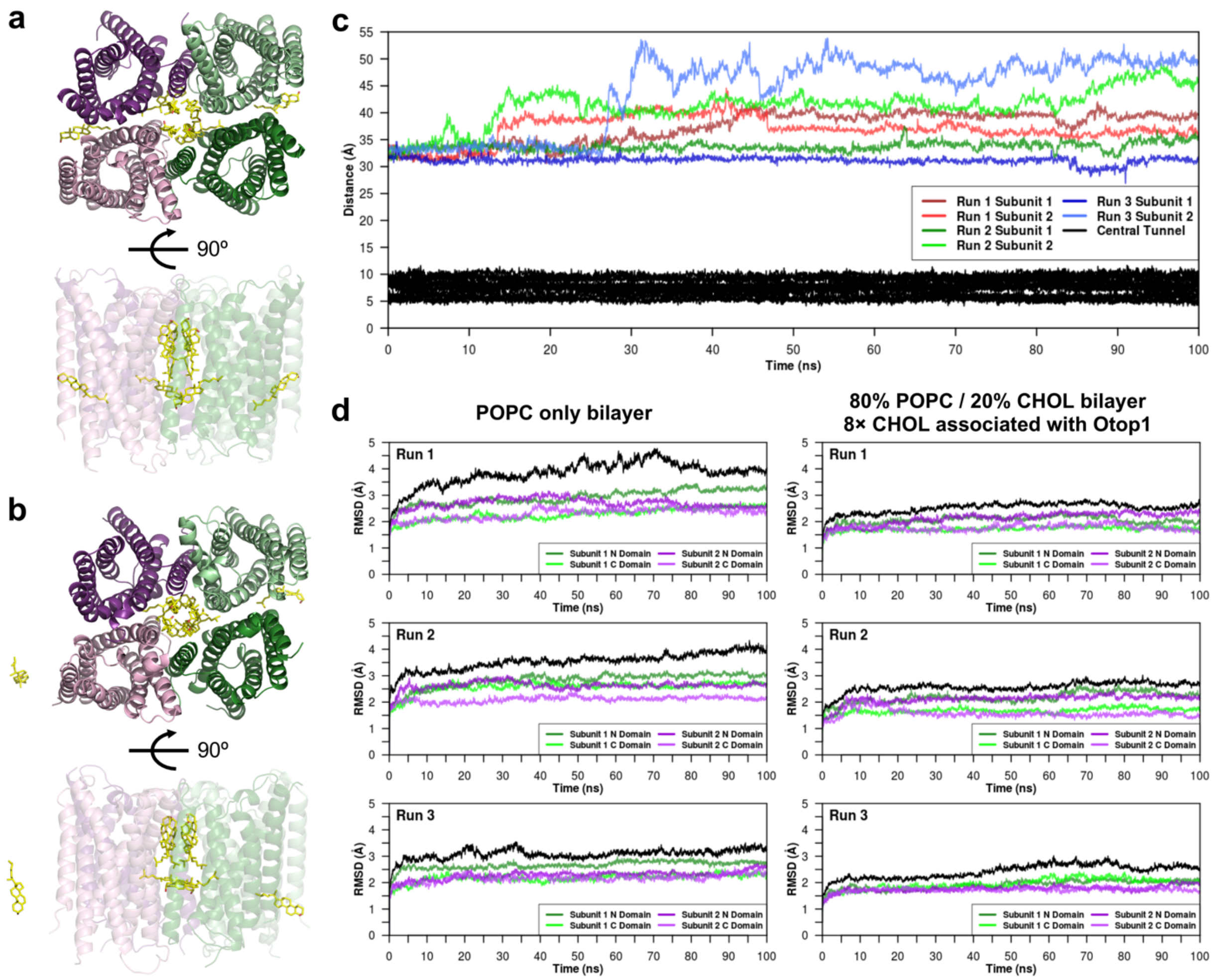


**Supplementary Figure 8. Dynamic arrangements of cholesterol molecules associated with Otop1 and their role in protein stabilization. a, b,** in MD simulations, the relative positions of the six cholesterol molecules occupying the central tunnel were maintained. In contrast, those cholesterol molecules occupying the intrasubunit interface have greater mobility. Snapshots from a representative run are shown here, corresponding to Run 3 in **c**. Coloring of the protein is consistent with Fig. 1. The upper and lower panels in **a** are top and side views at the start of the simulation, while those in **b** are the corresponding snapshots after the 100-ns all-atom simulation. **c,** distances between the center of geometry (COG) of Otop1 and the COGs of cholesterol molecules initially associated with the intrasubunit interface, plotted against simulation time (red, green and blue lines). Only in one of the six instances (Run 3, Subunit 1) the cholesterol molecule remained sandwiched between N and C domains throughout the 100 ns-simulation. The corresponding distances for the cholesterol molecules in the central tunnel are plotted as black lines, reflecting their stable arrangement within the tunnel. **d,** conformational drift of the protein is reduced in the presence of cholesterol molecules in the central tunnel and at the intrasubunit interface. Root-mean-square deviations (RMSDs) of the protein C-α atoms overall (black line) and at individual domain level (colors consistent with Fig. 1), with different membrane lipid compositions as indicated. Only residues of the TM helices were included in the calculation of RMSDs.

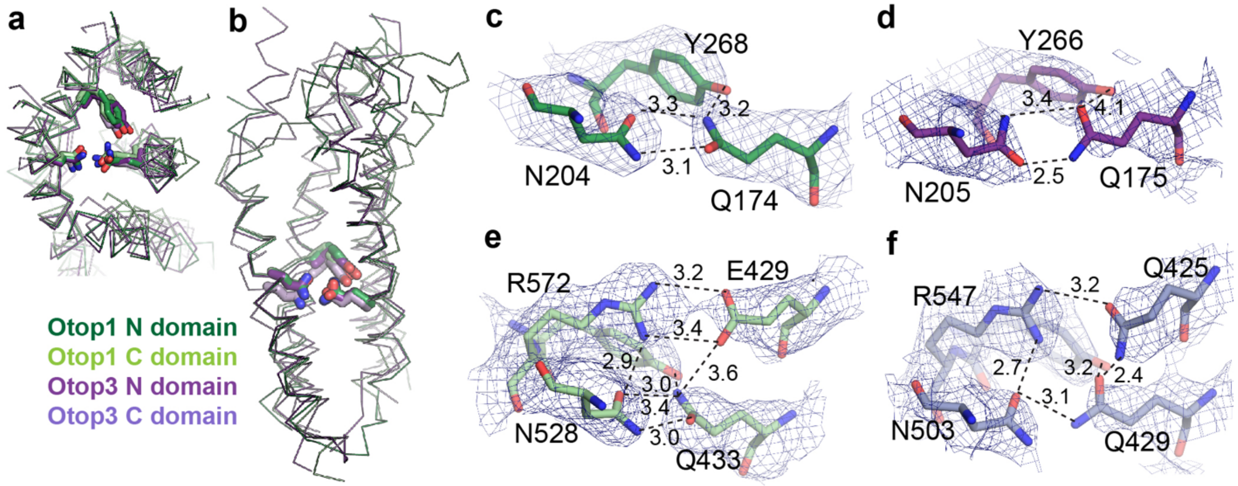


**Supplementary Figure 9. Conserved polar interactions in the putative Otop pores. a, b**, Top (**a**) and side (**b**) views of N domains and C domains of Otop1 and Otop3 aligned and superposed. TM1/7 are removed from **a** and TM1/2/7/8 are removed from **b** for clarity. The conserved QNY triad is shown as sticks. **c, d,** side view of the QNY triad in the N domains of Otop1 (**c**) and Otop3 (**d**). **e, f,** side view of the QNY triad in the C domains of Otop1 (**e**) and Otop3 (**f**), as well as additional interactions involving a conserved arginine and opposing glutamic acid (**e**) or glutamine (**f**). In **c-f,** sharpened cryo-EM maps (blue mesh, 5σ) are superimposed onto the models, and interatomic distances highlighting potential interactions are depicted with dashed lines and labeled with distances (in Å).

**a** N domain

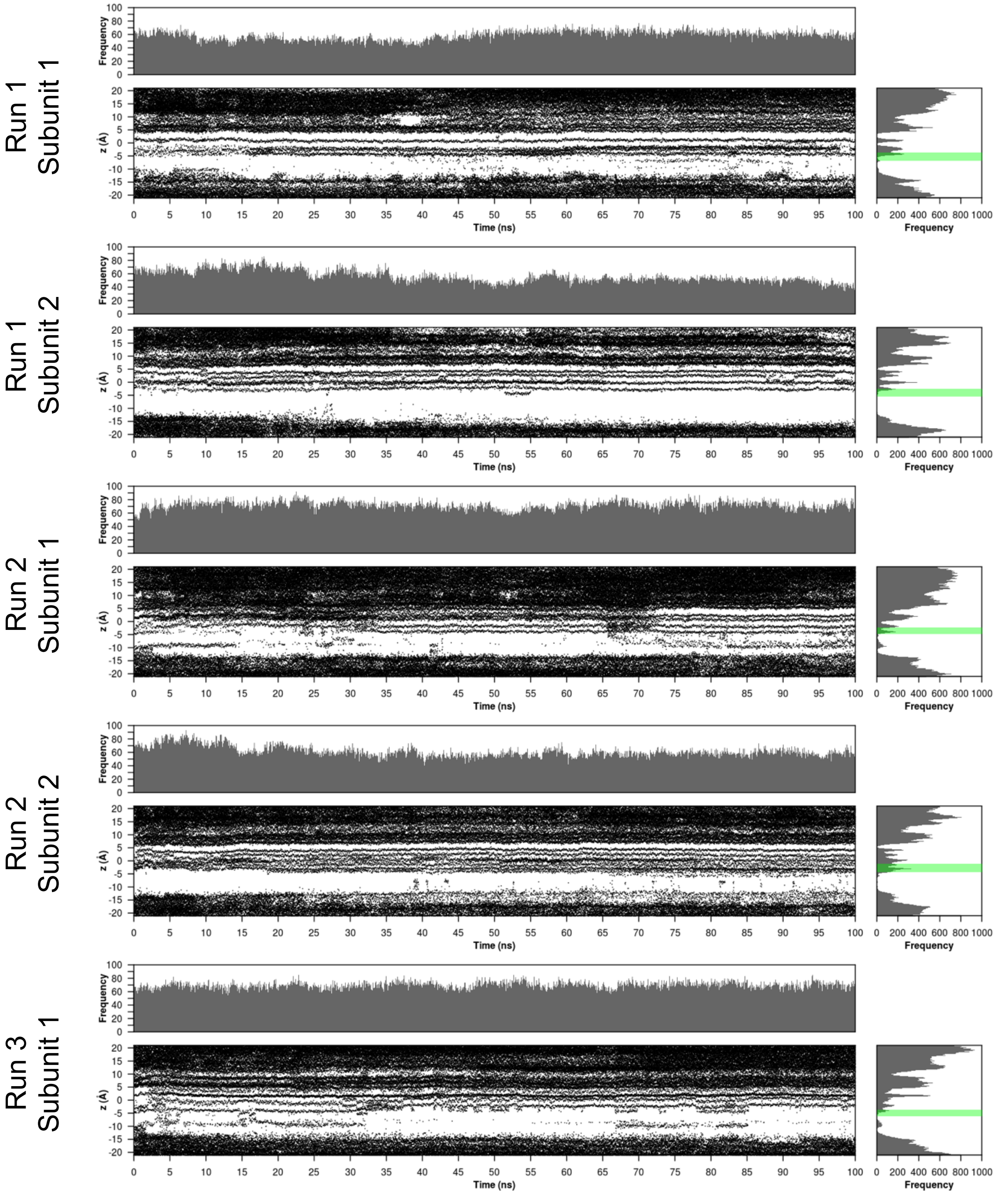

Run 1 Subunit 1
Run 1 Subunit 2
Run 2 Subunit 1
Run 2 Subunit 2
Run 3 Subunit 1
Frequency
z (Å)
Time (ns)

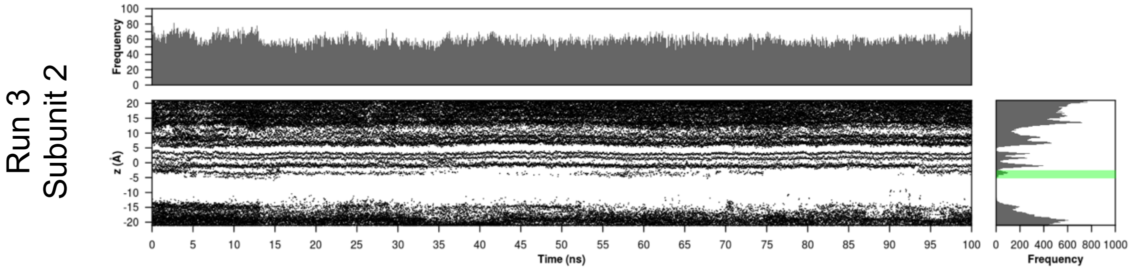
Run 3
Subunit 2
Frequency
z (Å)
Time (ns)
Frequency

**b** C domain

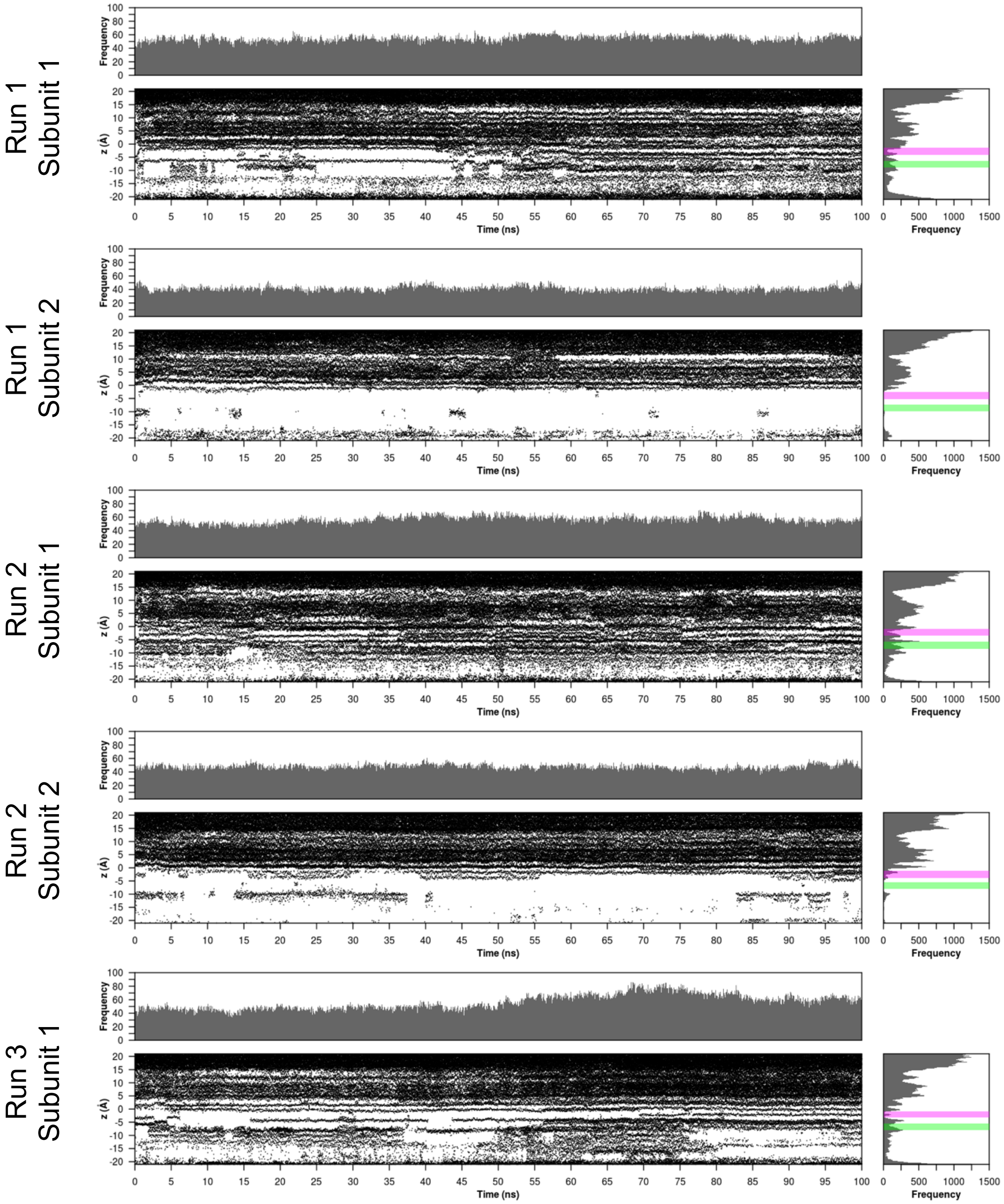
Run 1
Subunit 1
Run 1
Subunit 2
Run 2
Subunit 1
Run 2
Subunit 2
Run 3
Subunit 1
Frequency
z (Å)
Time (ns)
Frequency

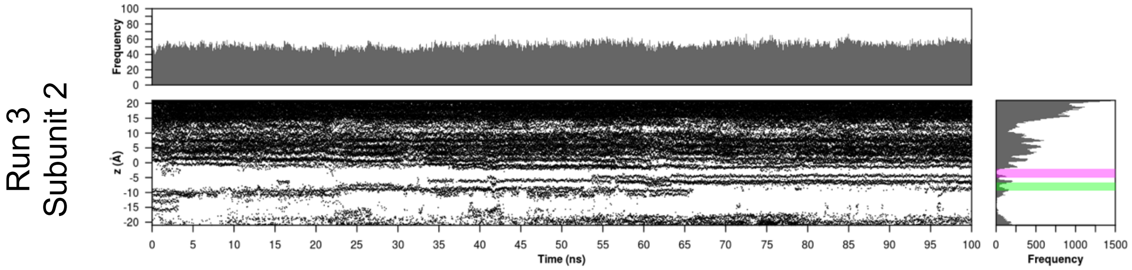


### c Intrasubunit Interface

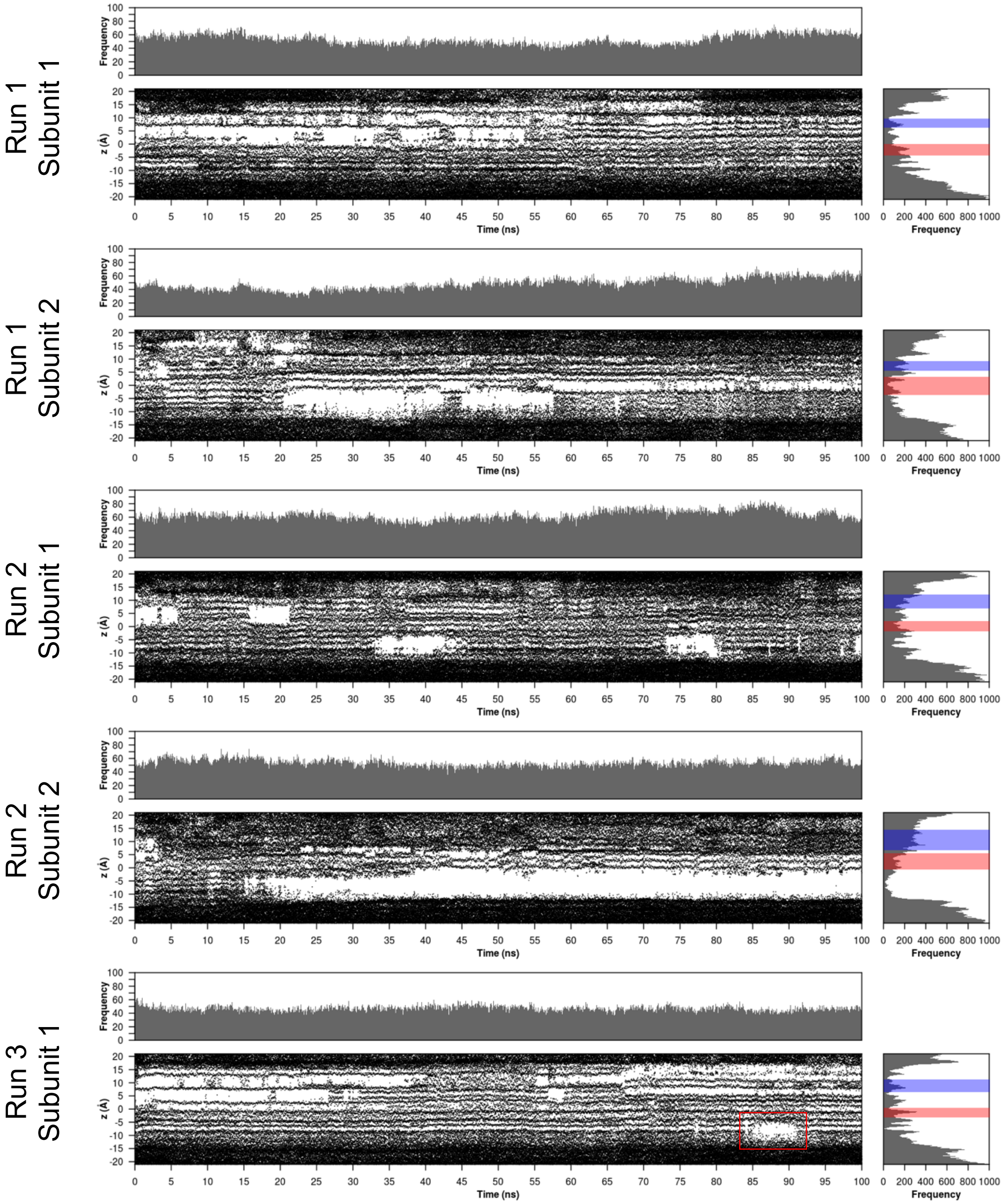

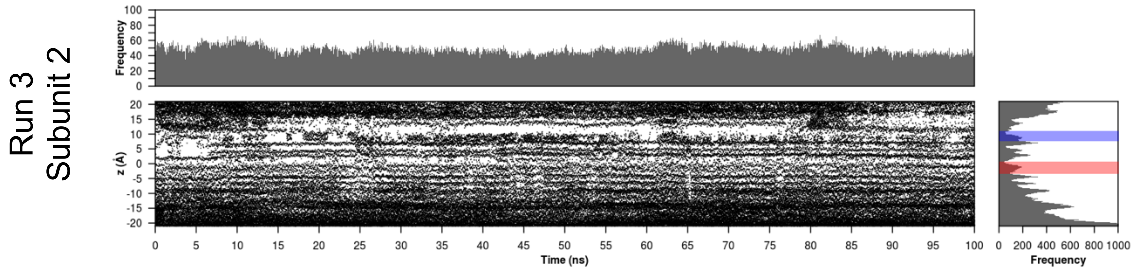


**Supplementary Figure 10. Trajectories of water oxygens along the three potential proton pathways, displayed for individual MD simulation runs and subunits. a,** data for the N domain. The region below the level of the QNY triad (green box in the right histogram) is stochastically wet and dewetted. **b,** Data for the C domain. Similar stochastic wetting and dewetting under the QNY triad is observed. The magenta box shows the position of the salt bridge formed by E429 and R572. **c**, Data for the intrasubunit interface. Overall, the distribution of water here is more uniform than that in the N and C domains. In 5 out of 6 instances the interface remains largely wetted in the simulation. The positions of R540 and E267 are denoted by the blue and red boxes in the right histogram. In the boxed region of Run 3 Subunit 1, the dewetting is concomitant with the transient insertion of the cholesterol tail (Supplementary Fig. 8c). Fig. 5g is obtained by summing the right-hand histogram in (**a-c**).

| | Otop1 in nanodiscs EMDB 9360 PDB 6NF4 | Otop3 in nanodiscs EMDB 9361 PDB 6NF6 |
|---|---|---|
| **Data collection and processing** | | |
| Magnification | 29,000 | 29,000 |
| Voltage (kV) | 300 | 300 |
| Electron exposure ($e^{-}/Å^2$) | 50 | 50 |
| Defocus range (μm) | 0.6-1.8 | 0.6-1.8 |
| Pixel size (Å) | 1.03 | 1.03 |
| Symmetry imposed | C2 | C2 |
| Initial particle images (no.) | 1,517,125 | 1,386,058 |
| Final particle images (no.) | 67,425 | 43,667 |
| Map resolution (Å) | 2.98 | 3.32 |
| FSC threshold | 0.143 | 0.143 |
| **Refinement** | | |
| Map sharpening *B* factor ($Å^2$) | -74 | -50 |
| Model composition (# of atoms) | | |
| Protein | 6694 | 6528 |
| Ligands | 266 | 70 |
| R.m.s. deviations | | |
| Bond lengths (Å) | 0.01 | 0.02 |
| Bond angles (°) | 1.00 | 0.95 |
| Validation | | |
| MolProbity score | 1.00 | 1.56 |
| Clashscore | 1.08 | 4.54 |
| EMRinger score | 3.04 | 2.76 |
| Poor rotamers (%) | 0.27 | 0 |
| Ramachandran plot | | |
| Favored (%) | 96.8 | 95.2 |
| Allowed (%) | 3.2 | 4.5 |
| Disallowed (%) | 0 | 0.3 |

**Supplementary Table 1. Statistics for data collection, data processing, model refinement, and validation.**
